# Eukaryotic-like gephyrin and cognate membrane receptor coordinate corynebacterial cell division and polar elongation

**DOI:** 10.1101/2023.02.01.526586

**Authors:** M. Martinez, J. Petit, A. Leyva, A. Sogues, D. Megrian, A. Rodriguez, Q. Gaday, M. Ben Assaya, M. Portela, A. Haouz, A. Ducret, C. Grangeasse, P. M. Alzari, R. Durán, A. Wehenkel

**Affiliations:** Structural Microbiology Unit, Institut Pasteur, CNRS UMR 3528, Université Paris Cité, F-75015 Paris, France.; Analytical Biochemistry and Proteomics Unit, Institut Pasteur de Montevideo, Instituto de Investigaciones Biológicas Clemente Estable, Montevideo, Uruguay.; Plate-forme de cristallographie, C2RT-Institut Pasteur, CNRS, UMR 3528, Université Paris Cité, F-75015 Paris, France.; Molecular Microbiology and Structural Biochemistry, CNRS UMR 5086, Université de Lyon, 7, passage du Vercors, 69367 Lyon, France.

## Abstract

The order *Corynebacteriales* includes major industrial and pathogenic actinobacteria such as *Corynebacterium glutamicum* or *Mycobacterium tuberculosis*. Their elaborate multi-layered cell wall, composed primarily of the mycolyl-arabinogalactan-peptidoglycan complex, and their polar growth mode impose a stringent coordination between the septal divisome, organized around the tubulin-like protein FtsZ, and the polar elongasome, assembled around the tropomyosin-like protein Wag31. Here, we report the identification of two new divisome members, a gephyrin-like repurposed molybdotransferase (GLP) and its membrane receptor (GLPR). We show that the interplay between the GLPR/GLP module, FtsZ and Wag31 is crucial for orchestrating cell cycle progression. Our results provide a detailed molecular understanding of the crosstalk between two essential machineries, the divisome and elongasome, and reveal that *Corynebacteriales* have evolved a protein scaffold to control cell division and morphogenesis similar to the gephyrin/GlyR system that in higher eukaryotes mediates synaptic signaling through network organization of membrane receptors and the microtubule cytoskeleton.

## INTRODUCTION

Cell division is central to bacterial physiology. The fundamental discovery by Francois Jacob in 1968 of the filamentation temperature-sensitive (fts) genes in *E. coli* led to the discovery of the tubulin like bacterial cytoskeleton protein FtsZ ^1^. Since then, and thanks to major technical developments in cell imaging and structural biology, well studied model systems set the basis for our current knowledge at the molecular level. While many cell division genes and interaction networks were identified in these model systems, the detailed molecular mechanisms underlying bacterial cell division remain enigmatic, notably because of the diversity of mechanisms specific to the different bacterial physiologies ^2–4^. FtsZ is a hub for protein-protein interactions that, through GTP-dependent polymerization, scaffolds and regulates the assembly of the cell division machinery (the divisome) at the site of septation and governs the ordered assembly of the cell wall biosynthetic machinery ^4^. In actinobacteria, a large and ancient phylum that includes important human pathogens such as *Mycobacterium tuberculosis* or *Corynebacterium diphtheriae*, many of the well-studied components of the divisome in model species like *E. coli* and *B. subtilis*, are missing from the genomes ^5^. This is notably the case of proteins that directly interact with FtsZ, including FtsA, EzrA and ZipA and it remains an open question whether FtsZ is regulated similarly via many different interactors as seen in other species. To date, only the essential membrane anchor SepF has been unequivocally identified as a direct interactor of the C-terminal domain of FtsZ ^6, 7^. The apparent lack of divisome components is particularly intriguing, not only because *Corynebacteriales* are polar growing bacteria that need to coordinate the mid-cell division and elongation machineries at a precise moment of the cell cycle when the septum becomes a new pole ^6, 7^, but also because the divisome and the elongasome must cooperatively work together to build and remodel the complex multi-layer cell wall formed by peptidoglycan, arabinogalactan and the mycolate outer membrane ^8, 9^ before the final “V-snapping” step of cytokinesis ^10^.

How the polar elongasome is assembled and controlled remains to be elucidated. The actin-like MreB scaffold ^11, 12^ is absent from the genomes. Instead, the cytoskeletal DivIVA homologue Wag31, a dimeric tropomyosin-like coiled-coil scaffolding protein with an N-terminal membrane binding domain, is essential for the polar elongasome assembly. Wag31 is required for the rod-shaped morphology of *Corynebacteriales* ^13–15^, and is also involved in chromosome segregation. Several proteins have been described as putative Wag31 interactors, mainly from genetic and cellular experiments, such as the non-conserved ParA/B ^16, 17^, CwsA ^18^, RodA ^14^, or yet MksG ^19^, but direct *in vitro* evidence for direct protein-protein interactions is still missing. Wag31 has a subpolar localization and, concomitant with or soon after septum formation, migrates to the cell division site at mid-cell to eventually start assembling the daughter cell elongasome at the new cell pole ^20^. The old pole grows faster than the new pole, suggesting that full maturation of polar and subpolar assemblies occurs over time and is divisome independent ^21, 22^. To date there is no knowledge of how the division and elongation processes are related in space and time and to what extent Wag31 is directly involved in protein-protein interactions with other components of the cell division and elongation machineries.

In this work, we have discovered two new members of the corynebacterial divisome and dissected a regulatory network of the cell cycle that directly links FtsZ to Wag31. We show that this link is mediated by a gephyrin-like protein (GLP) and its membrane receptor (GLPR). Mammalian gephyrin is a moonlighting protein that plays an essential role in synaptic signaling, *via* clustering of the glycine receptor GlyR, and is also involved in molybdenum cofactor (MoCo) biosynthesis. Like gephyrin, GLP has undergone evolutionary repurposing, as its 3D structure is closely related to that of molybdopterin molybdotransferase MoeA, but the FtsZ-binding capacity specifically evolved in Actinobacteria. Our biochemical, structural and cellular studies show that GLP and GLPR form a tight complex that is part of the early divisome, where GLP directly binds FtsZ, and interferes with cell elongation via direct interactions of GLPR with Wag31, thus placing the gephyrin-like GLP-GLPR complex at the center of the divisome-elongasome transition. This finding paves the way towards the understanding of the interplay between cell constriction and elongation leading to two identical daughter cells.

## RESULTS

### Identification of a novel divisome component in Actinobacteria

To discover missing players in corynebacterial cell division we used extensive mass-spectrometry based interactomics, starting with the FtsZ membrane anchor SepF as bait for co-immunoprecipitation (co-IP) of divisome members. *Corynebacterium glutamicum* (*Cglu*) cultures were cross-linked during exponential growth to stabilize highly dynamic interactions or interactions that depend on spatial cues such as the inner membrane and the FtsZ polymerization status. We carried out the co-IPs using the fluorescent protein tag, mScarlet, as bait in *Cglu* strains expressing either SepF-mScarlet or mScarlet, as well as using anti-SepF antibodies in both the untransformed *Cglu* (ATCC13032) and the SepF-mScarlet strain. 20 proteins were exclusively detected or statistically enriched from *Cglu* when compared with the control (*Cglu*-mScarlet). Using the same criteria as above, 22 and 98 proteins were recovered from the *Cglu*_SepF-mScarlet strain using anti-SepF and anti-mScarlet antibodies respectively (Figure S1a and Table S1a). 11 proteins with a quantifiable enrichment factor (common to all IPs) represent the SepF core interactome (Figure 1a). This core interactome includes FtsZ as expected, but the most enriched protein compared to the *Cglu* proteome is Cgl0883. This top interactor appeared consistently in all replicates and is hereafter named GLP (explained below). In *Cglu*, GLP is annotated as one of three molybdopterin molybdotransferase MoeA (EC 2.10.1.1) enzymes produced by this organism. MoeA enzymes incorporate the molybdenum metal into the molybdopterin (MPT) precursor to form the Moco cofactor used by molybdoenzymes to catalyze redox reactions ^23^.

**Figure 1:**
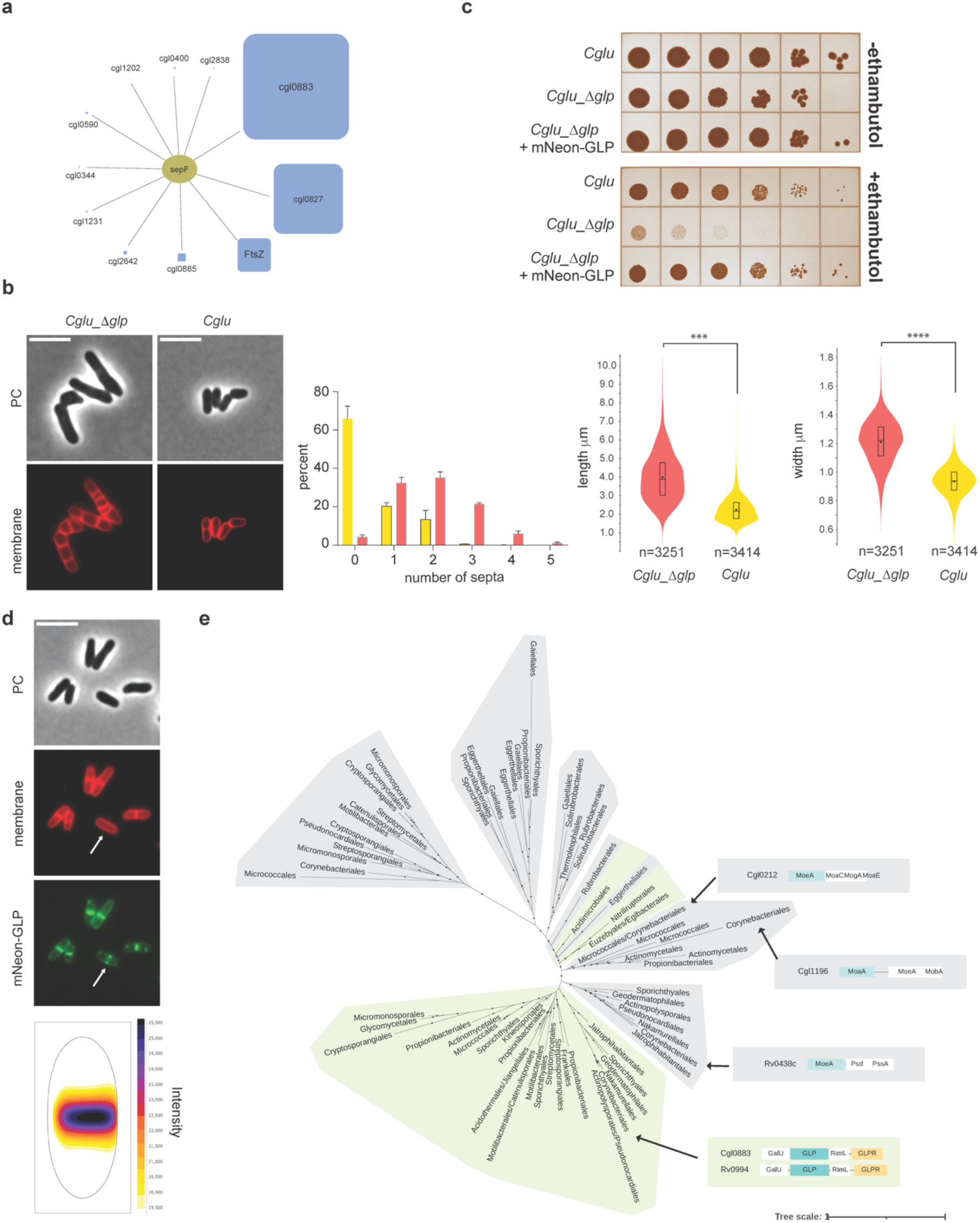
Identification of GLP as a new member of the corynebacterial divisome. (**a**) The core interactome of SepF, including proteins recovered from 3 independent co-IP experiments using different strains/antibodies: *Cglu*/*α*-SepF; *Cglu*_SepF-Scarlet/*α*-SepF and *Cglu*_SepF-Scarlet/*α*-Scarlet; performing for each one pairwise comparisons with controls. (Table S1 and Figure S1a). The square size for each interactor is proportional to its enrichment in the interactome compared to the proteome (Table S1). (**b**) GLP depletion. Representative images in phase contrast (upper row) and membrane staining (nile red) fluorescent signal (lower row) of *Cglu_Δglp* and *Cglu*. Frequency histogram indicating the number of septa per cell for *Cglu* (yellow) and *Cglu_Δglp* (red), calculated from n cells imaged from 3 independent experiments for each strain (*Cglu*, n=718, 1468 and 1223; *Cglu_Δglp,* n=873, 1538 and 840); bars represent the mean ± SD. Violin plots showing the distribution of cell length (***, d = 1,76, p ∼ 0) and cell width (****, d = 2,21, p ∼ 0) for *Cglu* (in yellow) and *Cglu_Δglp* (in red); the number of cells used (n) is indicated below each violin plot (cells from triplicate experiments); the box indicates the 25^th^ to the 75^th^ percentile, the mean and the median are indicated with a dot and a line in the box, respectively. (**c**) Ethambutol sensitivity assay. BHI overnight cultures of *Cglu* and *Cglu_Δglp* complemented with the empty plasmid or mNeon-GLP were normalized to an OD_600_ of 0.5, serially diluted 10-fold, and spotted onto BHI agar medium with or without 1 μg/ml (**d**) Localization of mNeon-GLP in *Cglu*. Representative images in phase contrast, membrane staining and mNeon-GLP fluorescent signal for the *Cglu*. The arrow indicates the GLP localization prior to septum formation. Heatmap representing the localization pattern of mNeon-GLP; 3879 cells were analyzed, from triplicate experiments. Scale bars 5μm. (**e**) Maximum likelihood phylogeny of MoeA paralogs in *Actinobacteria*. Clades with a green background correspond to GLP, while clades in gray correspond to other MoeA paralogs. The genomic context of GLP/MoeA is indicated for each gene present in *Cglu* (Cgl locus tag) and *M. tuberculosis* (Rv locus tag) genomes. Monophyletic classes were collapsed into a single branch for clarity. Dots indicate UFB > 0.85. The scale bar represents the average number of substitutions per site. For the detailed tree, see Supporting Data.

A GLP knock-out strain (*Cglu*_*Δglp,* Figure S1b) was viable but displayed a strong cell division phenotype, with elongated cells and multiple septa (Figure 1b), suggesting a delay in the final steps of cell division. The *Cglu*_*Δglp* strain was also sensitive to the anti-tuberculosis drug ethambutol (Figure 1c), a further indication of GLP involvement in cell division, as sub-lethal concentrations of ethambutol have been successfully used to identify genes required for cell growth and division ^24^. Both the multi-septate and ethambutol-sensitive phenotypes were restored to wild-type (WT) when GLP or mNeon-GLP were expressed from a plasmid under the control of P*_gntK_*, a tight promoter that is repressed by sucrose and induced by gluconate ^25^ (Figures 1c, S1b and S1c). Fluorescently labelled mNeon-GLP localized to mid-cell prior to septum formation, placing it with the early arrivers to the site of cell division (Figure 1d). In contrast, the two paralogs of GLP in the *Cglu* genome, MoeA1/Cgl0212, (25% aa sequence identity) and MoeA3/Cgl1196 (27% aa sequence identity), displayed a cytoplasmic distribution showing neither specific localization patterns nor morphological alterations when expressed as N-terminally tagged proteins with mNeon (Figure S1d). Their diffuse cytosolic signal was not due to fusion protein degradation (Figure S1e), suggesting that only GLP evolved specific functions related to the divisome.

While we identified MoeA paralogs in most Bacteria (Figure S2a), GLP seems to be present exclusively in Actinobacteria (Figure S2b). The phylogeny of all MoeA paralogs in Actinobacteria (Figure 1e) suggests that GLP was acquired by the ancestor of all Actinobacteria or early during its diversification and was later inherited vertically by most of its members. Our results suggest that most MoeA paralogs in Actinobacteria were acquired by several duplication or horizontal gene transfer events within Actinobacteria (Figures 1e and S2b). The genomes of most other bacterial phyla contain one MoeA paralog, which presumably corresponds to the MoeA enzyme, as in *Escherichia coli*. Taken together, these results suggest that GLP is a molybdotransferase-related enzyme that has acquired novel functions in Actinobacteria as a result of divergent evolution, a trait highly reminiscent of mammalian gephyrin.

### GLP is a gephyrin-like protein that interacts with FtsZ

The eukaryotic gephyrin is described as a moonlighting enzyme originally identified as a glycine receptor-associated two-domain protein in neurons ^26–28^. It was later found that the E-domain of Gephyrin corresponds to MoeA and that it functions as Moco biosynthetic enzyme ^29, 30^. Full-length gephyrin, which has an additional MogA domain, acts as a scaffold through oligomerization of its N- and C-terminal domains, via trimerization and dimerization respectively ^31^. It is known to transiently cluster and stabilize glycine (Gly) and GABAA receptors at the synapses of the mammalian brain ^30, 32^. It is thus tempting to speculate that GLP could also form a protein network in bacteria upon association with cell division proteins. Since there is evidence for a physical linkage between Gephyrin, GlyR and microtubules ^27^, we sought to investigate whether GLP septum localization could be accounted for by a direct physical interaction with the bacterial tubulin homologue FtsZ. Our interactomics data were consistent with this hypothesis because, when using a SepF mutant unable to bind FtsZ as bait, we saw a significant decrease of GLP binding (Figure 2a and Table S1b), suggesting that the observed SepF-GLP interaction occurred via FtsZ. This was further confirmed *in vitro* with purified proteins. We could not detect direct binding between SepF and GLP, but we could measure a direct interaction between GLP and FtsZ with an apparent *Kd* of 4.7 μM as determined by biolayer interpherometry (BLI) studies (Figures 2b and S3a). The interaction was stronger with polymerized FtsZ in the presence of GTP (Figure S3b), suggesting that GLP may preferentially recognize the assembled FtsZ polymer at the divisome. The interaction is mediated at least in part by the conserved C-terminal domain of FtsZ (FtsZ_CTD_), a known hub for protein-protein interactions (Figure S3c).

**Figure 2:**
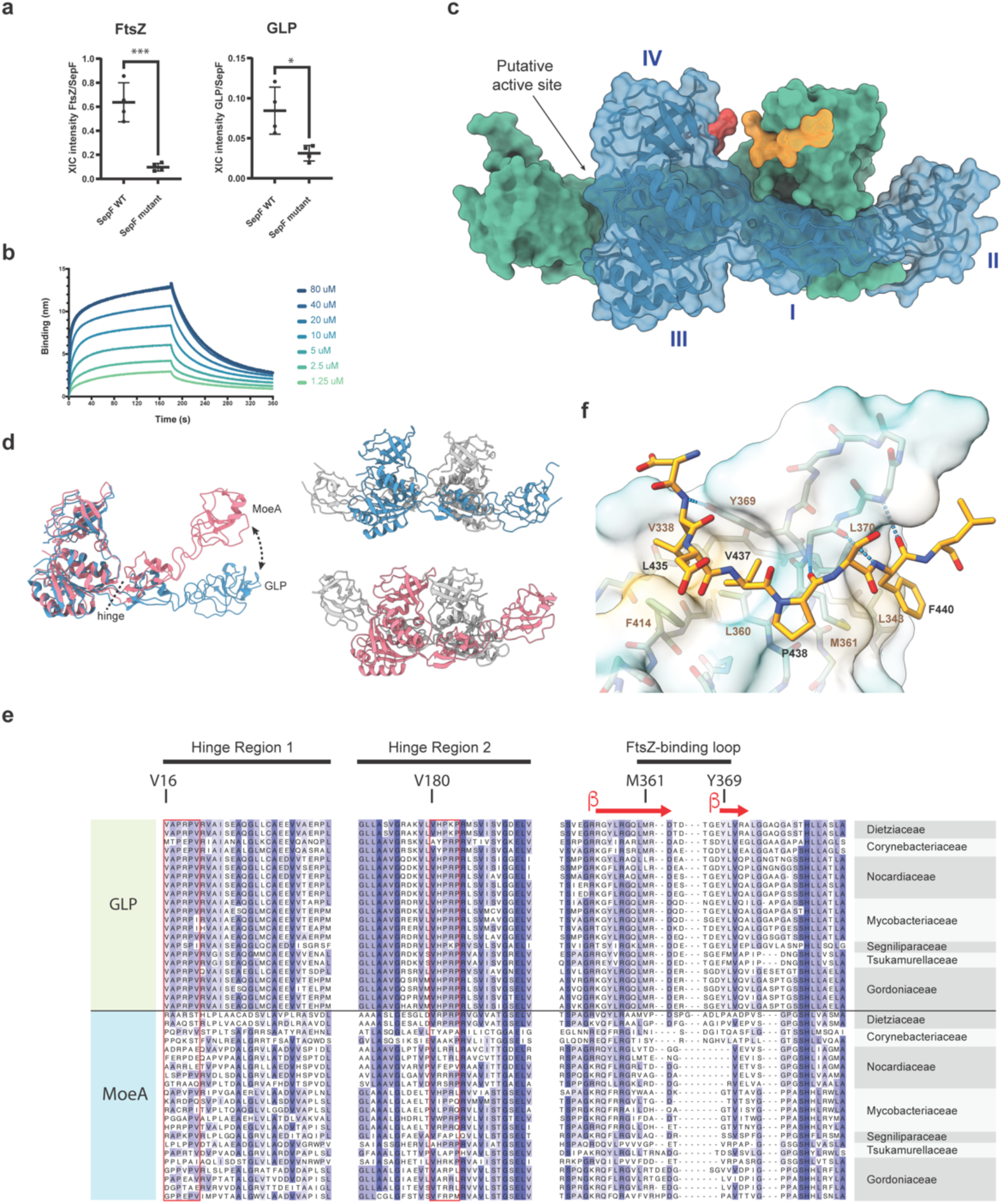
GLP-FtsZ interaction. (**a**) Comparison of the recovery of FtsZ and GLP in co-IP of *Cglu*_SepF-Scarlet-/*α*-Scarlet and the mutant unable to bind FtsZ (SepF_K125E/F131A_-Scarlet) using *α*-Scarlet. Each point corresponds to the normalized XIC intensity in each replicate of each condition, calculated as described in methods section; mean and SD are shown. Statistical analysis was performed using unpaired Student’s t-test (p < 0.05). FtsZ fold change = 6.61 (p value 0.0006); GLP fold change = 2.70 (p value 0.014). The corresponding analysis for each of the 11 core interactors is shown in Table S1b. (**b**) Sensorgrams of GLP binding to immobilized SUMO-FtsZ by biolayer interferometry. A series of measurements using a range of concentrations for GLP (inset) was carried out to derive the equilibrium dissociation constant (*K*_d_) (fitting shown in Figure S3a). (**c**) Crystal structure of the GLP homodimer (blue and green) in complex with FtsZ_CTD_ (yellow and red). The GLP monomer is composed of 4 structural domains (labelled I-IV in the blue monomer): domain I (residues 20-45 and 146-181), domain II (residues 46-145), domain III (residues 1-19 and 182-331), and domain IV (residues 332-417). The location of the putative active site at the distal dimer interface is also indicated. (**d**) Left panel: The superposition of the monomers from GLP (blue) and MoeA from *E. coli* (pink, pdb 1g8l) reveals a pronounced conformational change from a hinge region at the interface between domain I and III. This change leads to a central open (GLP, blue) or closed (MoeA, pink) conformation in the respective homodimers (right panel). (**e**) Partial alignment of three selected regions from MoeA paralogs in *Corynebacteriales*. Sequences of GLP and MoeA are shown for the same species, representatives of all *Corynebacteriales* families. The FtsZ-binding loop is delimited by the key residues methionine (M361) and tyrosine (Y369) indicated according to their position on the *Cglu* sequence. The Pro-rich hinge regions 1 and 2 are indicated by a red rectangle and the first residue inside the box is numbered and highlighted above. (**f**) Detailed view of FtsZ-GLP interactions. Residues involved in protein-protein interactions are labelled, the molecular surface of GLP is shown according to hydrophobicity (yellow=hydrophobic, green=hydrophilic), and intermolecular hydrogen bonds are shown as blue dotted lines.

We crystallized full-length GLP, both alone and in complex with the 10-residue peptide FtsZ_CTD_. The apo-structure was solved at 2.1 Å resolution using SAD techniques on a platinum derivative and the structure of the complex was solved at 2.7 Å resolution by molecular replacement using the apo-structure as a model (Table S2). The overall structure of the GLP dimer (Figure 2c) and the monomer organization into four structural domains (I-IV) are very similar to those described for *E. coli* MoeA ^33^ or the E-domain of mammalian gephyrin ^34^. A notable difference however is a pronounced hinge motion in the two sequence segments connecting the structural domains I and III, which leads to a more open GLP homodimer compared to bacterial MoeA (Figure 2d). This conformational change, which is not induced by FtsZ binding (Figure S3d), appears to be linked to the presence of a poly-Pro motif that is conserved in the two hinge regions of GLP homologs, but is missing in non-GLP MoeA (Figure 2e). This hinge motion generates the binding sites for FtsZ within the open GLP dimer interface. In the crystal structure, the FtsZ_CTD_ peptide binds the GLP homodimer with a 2:2 stoichiometry in its central region, far from the putative Mo-active site (Figure 2c). The FtsZ_CTD_ is well defined in the electron density (Figure S3e) and interacts primarily with a protruding *β*-hairpin in GLP domain IV (Figure 2e). The peptide adopts a linear extended conformation, with a central kink promoted by the presence of Pro438 (Figure 2f). The C-terminal half of the peptide backbone runs roughly parallel to the GLP *β*-strand 360-363 and is stabilized by three intermolecular hydrogen bonds between main-chain atoms (N_R362_-O_P438_, O_R362_-N_F440_ and N_A364_-O_F440_) and by hydrophobic interactions of FtsZ Phe440 with GLP residues Leu343, Met361 and Leu370 (Figure 2f). On the N-terminal half, the side chains of FtsZ residues Leu435 and Val437 are anchored in a hydrophobic pocket defined by GLP residues Val338, Leu360, Tyr369 and Phe414 (Figure 2f).

To validate the observed GLP-FtsZ interaction, we produced a deletion mutant of the entire FtsZ binding loop (Figure 2e) in GLP, replacing the sequence segment Met361-Leu370 by a single glycine residue (GLP_Δloop_). The purified mutant protein was correctly folded as shown by circular dichroism (Figure S3f) but as expected, it was unable to interact with FtsZ *in vitro* (Figure S3g). Complementation of the *Cglu*_*Δglp* strain with mNeon-GLP_Δloop_ failed to restore the wild-type morphology phenotype or the septal localization (Figure S3h and S3i), further stressing the physiological relevance of the crystallographic GLP-FtsZ complex. Both the two Pro-rich regions responsible for the hinge motion between GLP and MoeA and the FtsZ-binding loop (Figure 2e) can discriminate GLP from non-GLP MoeA homologs and therefore represent a molecular signature of the functional MoeA repurposing for specific divisome functions. Taken together, the above results demonstrate that GLP has evolved to bind FtsZ and is recruited to the division site by the direct interaction of domain IV with the conserved C-terminal domain of polymerized FtsZ.

### GLPR, a membrane receptor for GLP

To further investigate the GLP function we performed the reverse interactome, this time using GLP as bait. The list of proteins recovered as GLP interactors, using *Cglu*_*Δglp* as a negative control, are shown in Table S1c. Besides recovering both FtsZ and SepF as expected, among the top exclusive GLP interactors we identified Cgl0885, a membrane protein of unknown function (named hereafter GLPR for <u>GLP r</u>eceptor) that was already present among the top SepF interactors (Figure 1a and Table S1a). As observed for GLP, GLPR also seems to be specific to Actinobacteria as we could not identify homologs in any other bacterial phyla. When present, genes *glpr* and *glp* co-occur (Figures 3 and S4), suggesting a common evolutionary history. Both *glpr* and *glp* are part of the same operon in *C. glutamicum* ^35^ and synteny analysis in Actinobacteria revealed a well conserved genomic context around these genes when present (Figures 1e and S2a), suggesting a functional link.

**Figure 3:**
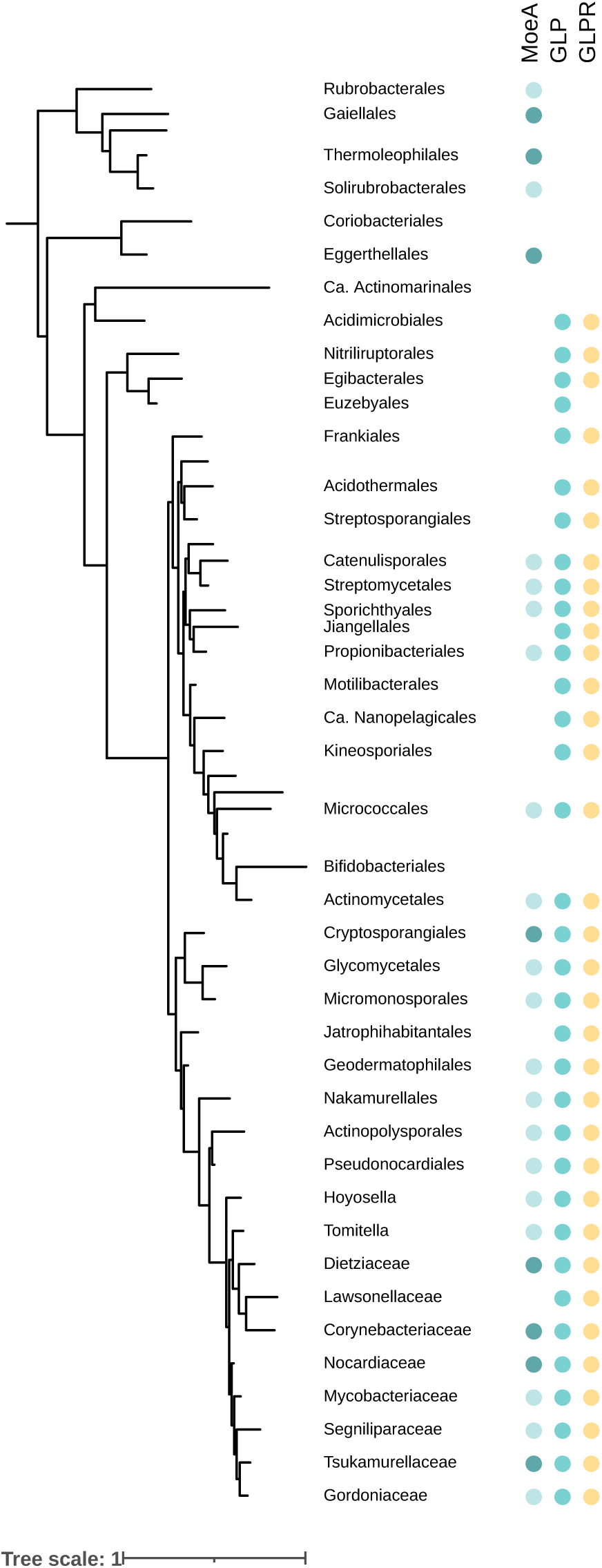
Phyletic pattern for the presence of MoeA, GLP and GLPR in Actinobacteria. Full circles indicate presence of the gene in more than 50% of the analyzed genomes of the phylum. Column MoeA indicates the presence of one (light blue) or more (dark blue) paralogs, except for GLP that is indicated in a separate column. The presence of GLPR is indicated by yellow dots. The phyletic pattern is represented on a reference *Actinobacteria* tree. *Actinobacteria* classes were collapsed into a single branch for clarity. Dots indicate UFB > 0.85. The scale bar represents the average number of substitutions per site. For the detailed tree see Figure S4, and for the detailed analysis see Table S3.

**Figure 4:**
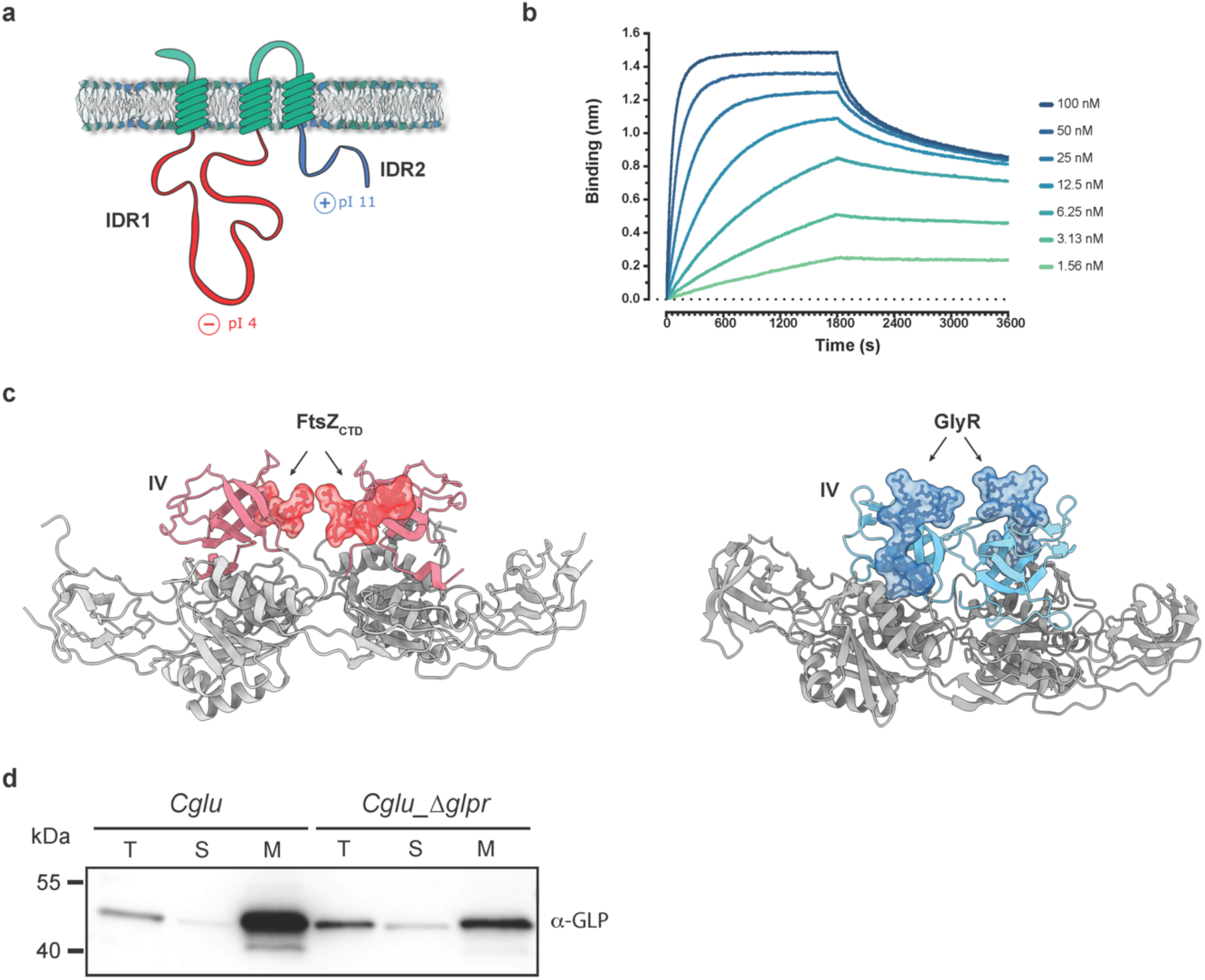
Identification of GLPR as a membrane receptor for GLP. (**a**) Schematic representation of GLPR, with the 3 transmembrane segments and the short external loops shown in green and the intrinsically disordered regions IDR1 (residues 27-218) and IDR2 (residues 263-340) in red and blue, respectively. The two IDRs are highly charged, with theoretical isoelectric points (pI) of 4.05 (IDR1) and 10. 87 (IDR2). (**b**) Sensorgrams of GLP binding to immobilized GLPR by biolayer interferometry. A series of measurements using a range of concentrations for GLP (inset) was carried out to derive the equilibrium dissociation constant (*K*_d_) (see Figure S5a). (**c**) Comparison of the GLP-FtsZ_CTD_ complex (red, left panel) with the Gephyrin-GlyR complex (blue, right panel, PDB code: 2fts). In both cases the FtsZ_CTD_ and GlyR peptides (molecular surfaces) bind the domain IV (in colour) of GLP and gephyrin respectively. (**d**) Cell fractionation and subcellular localization of GLP. Total (T), soluble (S) and membrane (M) fractions of *Cglu* or *Cglu*_*Δglpr* strains were obtained by differential centrifugation and analyzed by Western blot using an *α*-GLP antibody.

GLPR is an integral membrane protein with 3 predicted trans-membrane (TM) helices, and two cytoplasmic, oppositely charged intrinsically disordered regions (IDRs) (Figure 4a). To determine if there is a direct interaction between GLP and GLPR, we produced the recombinant proteins and assessed their interaction using biolayer interpherometry (BLI). Both proteins strongly interact with an apparent *Kd* of 5.5 nM (Figure 4b and S5a), indicating that GLPR might function as a membrane receptor for GLP. Deletion of the FtsZ-binding loop of GLP (GLP_Δloop_) also abolished the interaction with GLPR (Figure S5b), suggesting that both FtsZ and GLPR bind to an overlapping region at the center of the GLP dimer. Interestingly, the comparison of the known binding sites of GlyR on mammalian gephyrin and FtsZ_CTD_ (GLPR) on GLP showed that both interactions are mediated by the structural domain IV (Figure 4c). In cellular fractionation assays, GLP is found in the membrane fraction of the *Cglu* strain despite not having any membrane anchoring domains (Figure 4d). GLP localization to the membrane is reduced (though not abolished) in the *Cglu*_*Δglpr* depletion strain, suggesting the probable contribution of other proteins (for instance, FtsZ-SepF) to the membrane partitioning and septum localization of GLP. Taken together our results show that *Corynebacteriales* have evolved a gephyrin/GlyR-like system involved in cell division, prompting us to name the genes Cgl0883 as GLP (for Gephyrin-Like Protein) and Cgl0885 as GLPR (for GLP-Receptor).

### GLPR links the mid-cell divisome with the future polar elongasome

Unlike the *Cglu*_*Δglp* strain, depletion of GLPR in the *Cglu*_*Δglpr* strain (Figure S6a) did not show a significant morphological phenotype (Figure S6b) nor was it sensitive to ethambutol (Figure S6c). At low levels of exogenous expression in *Clgu* and *Cglu*_*Δglpr* cells (Figure S6d), GLPR-mNeon localized to the septum (Figure 5a), and the cells had a normal morphology (Figure 5b). However, and strikingly, higher levels of GLPR-mNeon expression (Figure S6e) led to a strong morphotype, which is characterized by the delocalization of the elongasome, as revealed by branching, that is aberrant pole formation along the lateral walls (Figure 5c). Accordingly, the cell projected surface area is significantly increased in *Cglu*_*Δglpr* + GLPR-mNeon (gluconate) when compared to *Cglu* (Figure 5d). The observed phenotype is likely due to steric hindrance induced by the presence of mNeon, as the untagged overexpression of full-length GLPR or GLPR lacking the C-terminal IDR (GLPR_ΔIDR2_) does not lead to branching (Figures 5c). In naturally branching actinomycetes such as *Streptomyces*, apical growth is directed by the essential coiled-coil protein DivIVA, which marks the hyphal site ^36, 37^. Similarly, Wag31, the *Corynebacteriales* homologue of *Streptomyces* DivIVA, specifically marks the sites of growth and its dysregulation results in polar growth from incorrect sites in *M. smegmatis* ^38, 39^, strongly suggesting that Wag31 delocalization is linked to the branching phenotype of the GLPR-mNeon overexpression strain (Figure 5c). In the multiseptated *Cglu_Δglp* strain we observed no branching but a clear accumulation of Wag31 at the different septa (Figure 5g), in some cases associated to noticeable rounding of the otherwise straight septal membrane (Figure 5g), reminiscent of premature pole formation.

**Figure 5:**
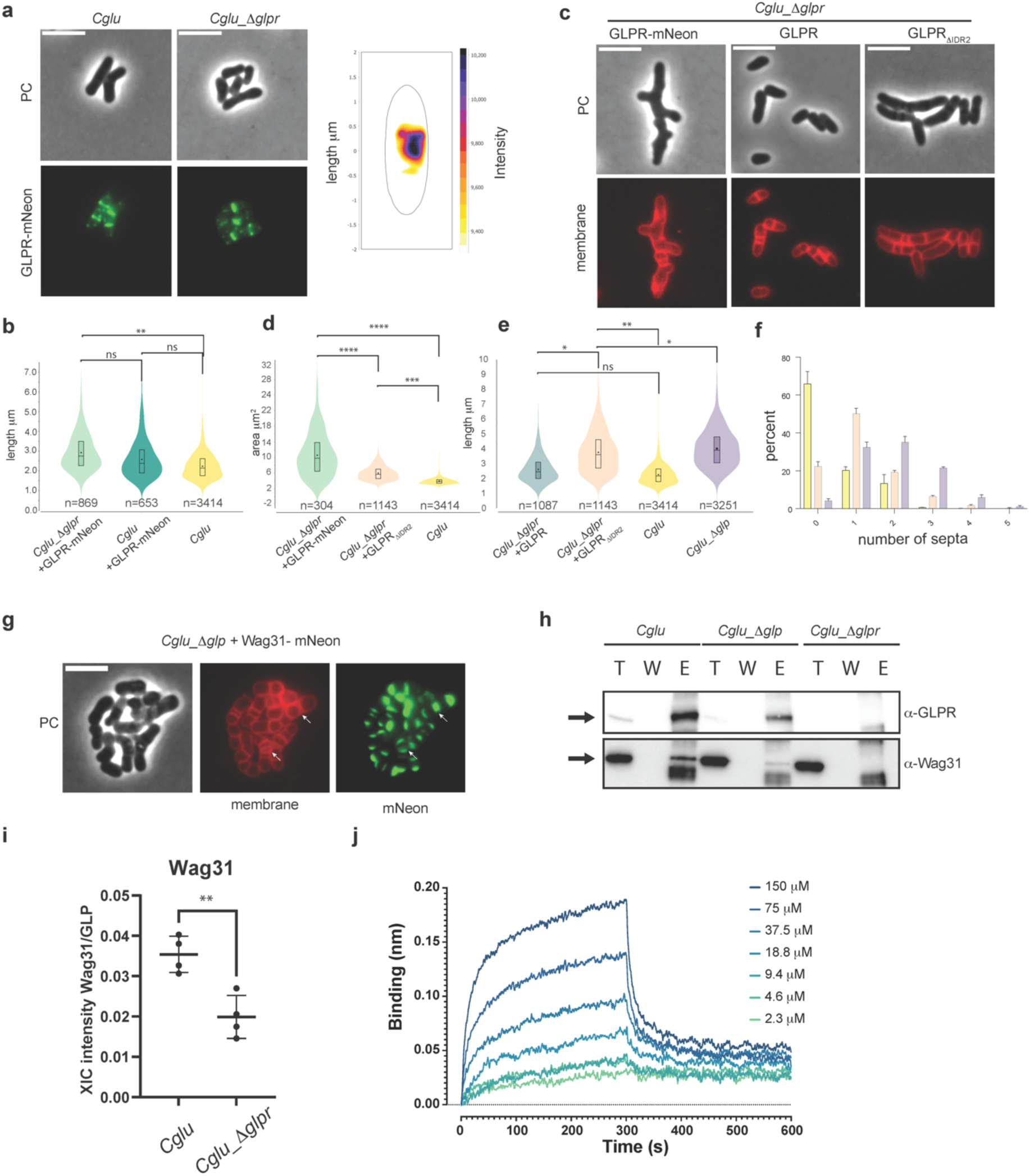
GLPR links the mid-cell divisome with the future polar elongasome *via* Wag31. (**a**) Representative images GLPR-mNeon expressed in *Cglu* and *Cglu_Δglpr*. The heatmap of the localization pattern of GLPR-mNeon in *Cglu*; 111 cells were analyzed. (**b**) Violin plots of cell length distribution for *Cglu_Δglpr* + GLPR-mNeon (light green), *Cglu* + GLPR-mNeon (dark green) and *Cglu* (yellow). Significance indicated corresponds to values of Cohen’s d: (**, d = 0,98, p = 2,88e-85), (ns, d = 0,49, p = 6,34e-19), (ns, d = 0,39, p = 2,78e-13)). The corresponding Western Blots are shown in Figure S6d. (**c**) Representative images of *Cglu_Δglpr* complemented with GLPR-mNeon, GLPR, or GLPR_ΔIDR2_ (in overexpression conditions of 1% gluconate). (**d**) Violin plots showing the distribution of cell surface areas for *Cglu_Δglpr* + GLPR-mNeon (green), *Cglu_Δglpr* + GLPR_ΔIDR2_ (orange) and *Cglu* (yellow). For *Cglu_Δglpr* + GLPR-mNeon, only cells showing a mean intensity of mNeon fluorescence greater than 35000 were considered, to discard cells that lost the plasmid. Significance Cohen’s d: (****, d = 3,95, p = 7,34e-63), (****, d = 2,35, p = 1,01e-46), (***, d = 1,4, p = 5,37e-319)). The Western blots of whole cell extracts of all those strains are shown in Figure S6e. (**e**) Violin plots of cell length distribution for *Cglu_Δglpr* + GLPR (blue), *Cglu* + GLPR_ΔIDR2_ (orange) and *Cglu* (yellow) and *Cglu_Δglp* (violet). Cohen’s d: (**, d = 1,17, p = 7,22e-255), (*, d = 0,61, p = 3,07e-60), (ns, d = 0,49, p = 8,12e-34), (*, d = 1,70, p = 6,24e-138, for *Cglu* + GLPR_ΔIDR2_ vs *Cglu_Δglp*)). (**f**) Frequency histogram of the number of septa per cell for *Cglu* (yellow), *Cglu_Δglp* + GLPR_ΔIDR2_ (orange) and *Cglu_Δglp* (violet), calculated from n cells from 3 independent experiments (*Cglu*, n=718, 1468 and 1223; *Cglu_Δglp* + GLPR_ΔIDR2_, n = 451, 737 and 841; *Cglu_Δglp,* n=873, 1538 and 840); bars represent the mean ± SD. For all violin plots: The box indicates the 25^th^ to the 75^th^ percentile, mean and median are indicated with a dot and a line in the box, respectively. The number of cells used in the analyses (n) below each violin representation corresponds to triplicates. (**g**) Localization of Wag31-mNeon in the multiseptated *Cglu*_*Δglp* strain (in overexpression conditions of 1% gluconate). Representative images are shown for phase contrast, Wag31-mNeon and nile red (membrane). The arrows indicate septum rounding inside the cell upon Wag31-mNeon overexpression in *Cglu*_*Δglp*. Scale bars 5μm. (**h**) Co-IP of GLPR-Wag31 in *Cglu*, *Cglu*_*Δglp* and *Cglu*_*Δglpr* strains. GLPR was used as bait. Total (T), wash (W) and elution (E) fractions were analyzed by Western blot using *α*-GLPR and *α*-Wag31 antibodies. Arrows indicate GLPR (top) and Wag31 (bottom). (**i**) Comparison of Wag31 recovery in co-IPs of GLP from *Cglu* and *Cglu_Δglpr* strains. Each point corresponds to the normalized XIC intensity in each replicate of each condition; mean and SD are shown. Statistical analysis was performed using unpaired Student’s t-test (p < 0.05). Wag31 fold change = 1.78 (p value 0.004). (**j**) Sensorgrams of Wag31 binding to immobilized GLPR by biolayer interferometry. A series of measurements using a range of concentrations for Wag31 was carried out to derive the equilibrium dissociation constant (*K*_d_) (Figure S7a).

The above results clearly demonstrate that interfering with GLP/GLPR interaction has pronounced effects on the regulation of the final steps of division (multisepta) and elongasome assembly and localization (branching) and suggest that this regulation may occur via excessive/premature Wag31 accumulation. More specifically, they point to a possible regulatory role of the GLP/GLPR complex on early elongasome assembly and localization, possibly acting on the elongasome scaffolding protein Wag31. This hypothesis was further confirmed by IP experiments showing complex formation *in vivo* between Wag31 and GLPR. Pulling on GLPR with an anti-GLPR antibody followed by Western Blot against the N-terminal DivIVA domain of Wag31 showed the co-elution of the two proteins in the wild-type strain (Figure 5h). This co-elution was reduced in *Cglu*_*Δglp* and was not seen in *Cglu*_*Δglpr*. Moreover, quantitative analysis of the MS experiments revealed not only that Wag31 was systematically enriched in the GLP interactome (Table S1c), but also that it was significantly decreased in the GLP interactome carried out in a *Cglu*_*Δglpr* background (Figure 5i). To seek direct biochemical evidence, we purified the three full-length proteins (GLP, GLPR and Wag31) as well as the N-terminal DivIVA-domain of Wag31 (Wag31_1-61_) for *in vitro* interaction studies. We were unable to detect any interaction between GLP and Wag31 (or Wag31_1-61_) under the conditions tested. In contrast, GLPR did bind both full-length Wag31 as well as Wag31_1-61_ with apparent *Kd* values of 43.4 μM and 14.9 μM, respectively, as measured by BLI (Figures 5j, S7a and S7b), demonstrating that Wag31-GLPR complex formation is mediated at least in part through the N-terminal DivIVA domain. Taken together, the above data demonstrate that the FtsZ-associated GLP/GLPR complex regulates early elongasome assembly at mid-cell *via* direct interaction with Wag31, the scaffolding protein of the elongasome.

## DISCUSSION

In this work we have identified GLP, a gephyrin-like repurposed molybdotransferase MoeA enzyme, and its membrane receptor, GLPR, as new components of the corynebacterial divisome. We show that GLP and GLPR are central elements of a protein-protein interaction network directly connecting the cytoskeletal proteins from the divisome (FtsZ) and the elongasome (Wag31) (Figure 6a). The structure of GLP is consistent with its annotation as MoeA, the enzyme involved in the synthesis of the Molybdenum cofactor (Moco), which is present in all forms of life and is used by molybdoenzymes to execute key transformations in the metabolism of nitrogen, sulfur and carbon compounds. Molybdoenzymes mediate essential cellular functions such as energy generation and detoxification reactions ^40^ and are believed to support virulence in pathogenic bacteria ^40, 41^. MoeA containing proteins have also acquired additional functions and act as moonlighting proteins ^30^. While this feature was thought to be a recent evolutionary trait restricted to eukaryotic Moco biosynthetic enzymes ^26^, the results presented here show that this is also the case for GLP in *Corynebacteriales*. Thus, the crystal structure of the GLP-FtsZ complex reveals the precise mode of binding of the FtsZ-CTD to GLP and provides a molecular signature for the evolutionary repurposing of the molybdotransferase. GLP has evolved a specific grove at the dimer interface that creates the FtsZ binding pocket for a 2:2 stoichiometric complex. Importantly, these results show that the corynebacterial FtsZ-CTD also acts as a hub for protein-protein interactions in complex and dynamic protein-protein association networks that govern cell division. The here identified network would make of FtsZ the eventual regulator of early elongasome assembly and maturation (Figure 6b, left panel), implying that the Z-ring cytoskeleton would ultimately be responsible for the septal localization of Wag31. Timely removal of this control (by Z-ring disassembly and cytokinesis) would lead to full maturation of the new pole linked to further Wag31 accumulation and late polar elongasome assembly (Figure 6b, right panel).

**Figure 6:**
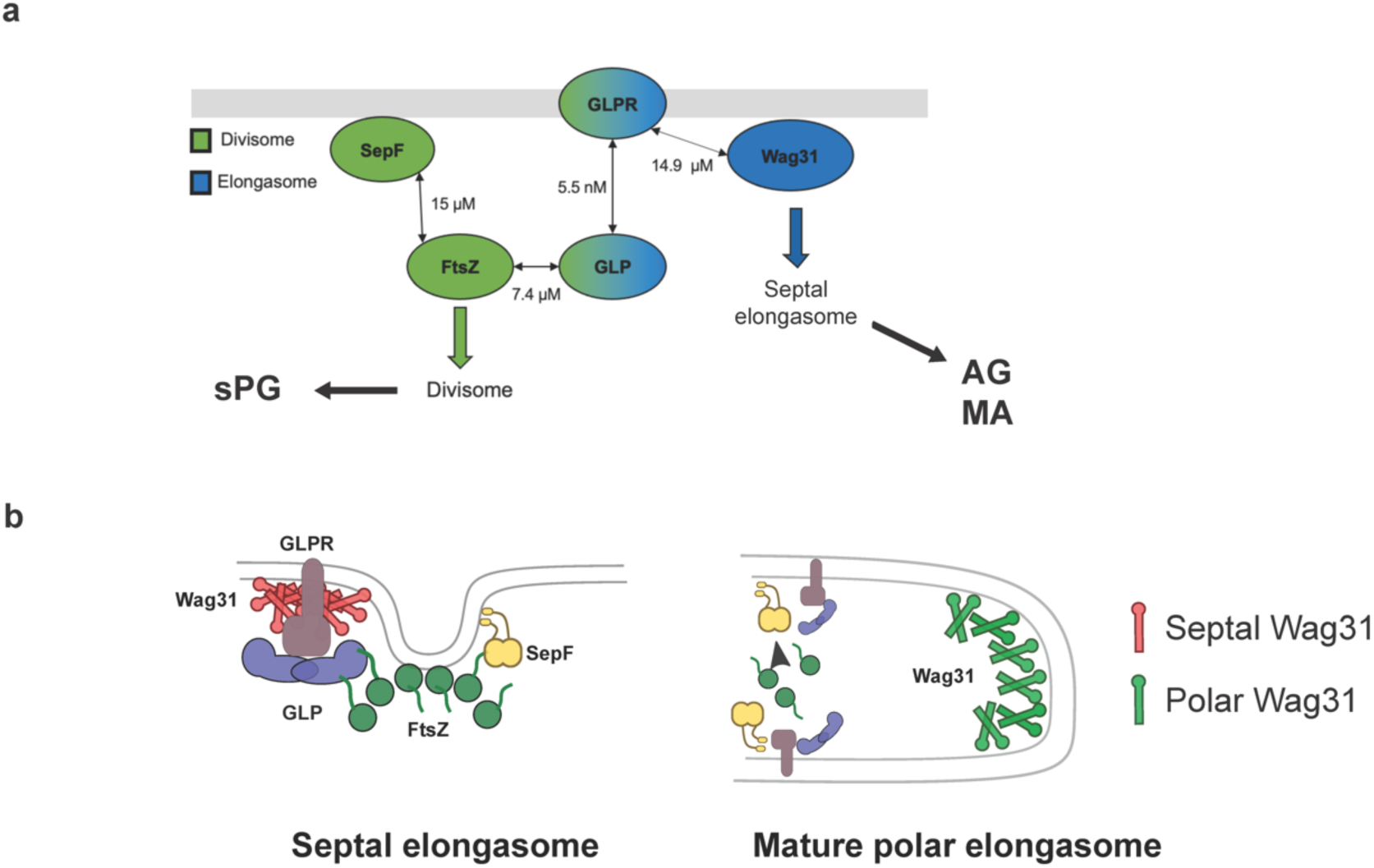
Interaction network and proposed function for GLP-GLPR. (**a**) The known direct interactions and their associated apparent K*_d_* values. Note that the SepF-FtsZ was determined using SPR (Sogues et al, 2020), whereas all other measurements were done with BLI (this work). (**b**) Working model on the roles of GLP-GLPR-Wag31 in the divisome-elongasome transition during cytokinesis in *Corynebacteriales*. At the septum, GLP-GLPR would control the functional status of Wag31 and prevent a premature pole formation through excessive Wag31 accumulation. Once cell division is completed, this septal control on Wag31 will disappear and an elongation competent cell pole can form.

Wag31 appears at mid-cell very early in the cell cycle and accumulates asymmetrically at the cell poles over time ^20^. This asymmetry is correlated with differential polar growth, the old pole growing faster than the new pole ^21, 22, 42^. The coiled-coil-rich Wag31 has a high propensity to self-associate and build higher order assemblies or networks (full-length Wag31 easily forms gels *in vitro*, see Materials and Methods) ^43^ and large foci of Wag31 are seen at the cell poles *in vivo*. As Wag31 arrives early to the septum, its self-association must be controlled to avoid premature pole formation. This negative regulation might depend on different aspects, such as conformational states of Wag31 and/or protein concentration and/or post-translational modifications (PTMs). Indeed, phosphorylation of Wag31 has been shown to be functionally important in *Mycobacteria* as well as *Streptomyces* ^15, 44^. Whichever the case, the initial control of Wag31 accumulation at mid-cell most likely depends on the divisome. Evidence for this comes from cellular studies where, upon conditional depletion of essential divisome components such as SepF, FtsQ and others ^7, 45, 46^, cells start branching (i.e., they assemble new poles in erroneous places over the lateral cell walls). Although Wag31 is most abundantly localized at the poles in WT cells, neither GLP not GLPR are found there (Figure 1d and 5a) except when Z-ring/divisome formation is abolished by depleting SepF (Figure S8). This is consistent with FtsZ retaining GLP/GLPR at the septum to specifically interact with and regulate Wag31, possibly in a coiled-coil conformation differing from the polar one, to preclude premature pole formation (as suggested by the septal membrane bending promoted by the absence of GLP, Figure 5g).

We observed erratic pole formation when overexpressing GLPR-mNeon (Figure 5c), indicating a functional interference of the mNeon tag on the FtsZ-GLP-GLPR-Wag31 network and subsequent Wag31 delocalization to induce polar growth from incorrect sites. However, removing GLP or the C-terminal IDR2 from GLPR results in an elongated multiseptal phenotype and septal elongasome dysregulation, but not branching (Figures 1b, 5c and 5e), probably because these deletions are less disruptive for network formation than the intercalation of the mNeon protein. Together, these observations suggest that interfering with the regulatory function of the GLP/GLPR complex leads to a delay in cell separation, possibly by dysfunction of the early septal elongasome that is required to synthetize the outer layers of the cell envelope during cell division ^10^. Early elongasome assembly starts while the divisome is still in place, as the two protein machineries are required for the synthesis of the full cell wall before cytokinesis. Two distinct enzyme systems exist to incorporate the septal and polar peptidoglycan (sPG and pPG), respectively orchestrated by the SEDS pair of enzymes FtsW/FtsI in the divisome and RodA/b-PBP in the polar elongasome. However, the synthesis and incorporation of the outer layers of the cell envelope is catalyzed by a unique set of enzymes, which belong to the elongasome but are also required to finalize cell separation ^10^. Evidence for this comes from experiments done with the anti-tuberculosis drug ethambutol, which target arabinosyltransferases (EmbA-C) and affects specifically the elongasome ^47^, but not sPG assembly and divisome function. The differential behavior of GLP and GLPR towards ethambutol (Figure S6c) and their respective interactions with scaffolding proteins (GLP-FtsZ and GLPR-Wag31) lends further support to the GLP/GLPR complex acting as a bridge between the late divisome and the early elongasome.

In *Bacillus subtilis* the Wag31 homologue, DivIVA recognizes negative membrane curvature, possibly through oligomerization into large networks (Lenarcic *et al*, 2009; Ramamurthi & Losick, 2009; Oliva *et al*, 2010). The DivIVA structural arrangement of the coiled-coil domain in *B. subtilis* ^48^ bears a resemblance to eukaryotic BAR domains ^49^, which bind to curved membranes but can also induce membrane deformation. Our results suggest that, besides recognizing negative membrane curvature ^50, 51^, Wag31 networks might contribute above all to define rod-like cell morphology by modulating polar membrane curvature once the septal constraints have been released (*i.e*., after daughter cell separation). Such a scenario would imply that Wag31 accumulation *per se* should be able to initiate a branching point along the lateral cell wall. This is indeed observed in mycobacterial coccoid cells, depleted for Wag31, where upon repletion Wag31 accumulation in a single spot induces a polar protrusion ^52^. Similarly, in *Streptomyces*, upon large polar accumulation of Wag31, small foci detach to assemble a new branching point (polarisome) away from the original Wag31 apical network^36^. Our cellular data show that in the absence of GLP and in the presence of excess Wag31, pole-like, rounded structures can form inside the cell, possibly driven by uncontrolled Wag31 accumulation (Figure 5g), suggesting a premature pole formation without timely completion of cell division.

In humans, gephyrin is an essential protein for clustering glycine (GlyR) and GABA receptors at the inhibitory synapse, a process thought to be mediated by the underlying tubulin and/or actin cytoskeletons ^53–56^. Although the gephyrin/GlyR and GLP/GLPR complexes are involved in unrelated biological processes (synaptic signaling and cell division), we can draw a molecular analogy between these two networks. They both have undergone evolutionary repurposing or acquired moonlighting functions from a common enzymatic scaffold (MoeA), and both interact with or organize the tubulin cytoskeleton. They are associated to the membrane through a tight complex formed between the soluble MoeA domain and their membrane receptor. In both cases the affinities are in the low micromolar or nanomolar range (this work; ^57–59^), contrasting with the more dynamic and lower affinity interactions with FtsZ or Wag31 for instance. Until now, repurposing of gephyrin was thought to be a trait reserved to recently evolved species such as *Homo sapiens* ^29^. At the light of our results on bacterial GLP, it is intriguing and remains an open question whether MoeA repurposing/moonlighting and its link to the tubulin cytoskeleton are inherited traits or evolutionary independent events. In any case, it appears that the MoeA scaffold has a propensity to acquire functions related to network formation and control at the inner membrane of cells in as crucially important processes as mammalian synaptic signalling and bacterial cell division.

## MATERIALS AND METHODS

### Bacterial strains, plasmids and growth conditions

All bacterial strains and plasmids used in this study are listed in the Table S5. *Escherichia coli* DH5α or CopyCutter EPI400 (Lucigen) was used for cloning and grown in Luria-Bertani (LB) broth or agar plates at 37°C supplemented with 50 µg/ml kanamycin when required. For protein production, *E. coli* BL21 (DE3) was grown in 2YT broth supplemented with autoinduction medium (0.5% glycerol, 0.05% glucose, 0.2% lactose) and 50 µg/ml kanamycin or 50 µg/ml carbenicillin at the appropriate temperature. *C.* glutamicum ATCC (*Cglu*) 13032 was used as a wild-type (WT) strain. *Cglu* strains were grown in brain heart infusion (BHI) or CGXII minimal medium ^60^ at 30°C and 120 rpm shaking supplemented with 25 µg/ml kanamycin when required. When specified, CGXII containing 4% sucrose was supplemented with 25 µg/ml kanamycine and/or 1% gluconate.

### Ethambutol (EMB) sensitivity assay

Overnight BHI cultures of *Cglu* and derivative strains were normalized to an OD_600_ of 0.5, serially diluted, and spotted (10 μl) onto BHI agar medium with and without 1 μg/ml EMB as indicated. Plates were incubated for 24 hours at 30°C and imaged using a ChemiDoc Imaging System (Bio-Rad).

### Cgl0883 (glp) and Cgl0885 (glpr) deletion in C. glutamicum

We used the two-step recombination strategy with the pk19mobsacB plasmid to delete the coding region of GLP. We amplified approximately 600 bp upstream and downstream of *glp* or *glpr* using chromosomal DNA of *Cglu* as a template. The PCR fragments were cloned by Gibson assembly into a linearized pk19mobsacB, obtaining the plasmid pk19-Δ*glp* or pk19-Δ*glpr*. The plasmids were sequence verified (Eurofins, France) and electroporated into *Cglu*. Insertion of the plasmids was checked by colony PCR and positive colonies were grown in BHI media supplemented with 25 μg/ml kanamycin overnight. The second round of recombination was selected by growth in BHI plates containing 10% (w/v) sucrose. Kanamycin sensitive colonies were screened by colony PCR to check for *glp* or *glpr* deletion. Positive colonies were sequence verified (Eurofins, France).

### Cloning for recombinant protein production in *E. coli*

Primers used for PCR amplification of the different fragments or site-directed mutagenesis are listed in Table S6. Cloning was performed by assembling the purified PCR fragments into the specified pET derivative expression vector using the commercially available NEBuilder HiFi DNA Assembly Cloning Kit (New England Biolabs).

The gene encoding for GLP was amplified by PCR using gDNA of *C. glutamicum* ATCC 13032 as template and cloned into a pET vector containing an N-terminal 6xHis-SUMO tag by Gibson assembly. GLP_ΔLoop_ (in which residues 362-371 were replaced by a single glycine residue) was generated by site-directed mutagenesis using the plasmid pET-SUMO-GLP as template.

Likewise, the gene coding for Wag31 was amplified by PCR using gDNA of *Cglu* as template and cloned into a pET vector containing an N-terminal 6xHis-SUMO tag by Gibson assembly. The plasmid encoding Wag31 with an N-terminal 6xHis tag followed by a TEV protease cleavage site (pET-His-TEV-Wag31) was synthesized by Genscript. The Wag31 N-terminal DivIVA domain (Wag31_1-61_) was generated from the pET-His-TEV-Wag31 plasmid by introducing a STOP codon after residue 61 by PCR mutagenesis.

The gene encoding for GLPR was amplified by PCR using gDNA of *Cglu* as template and cloned into a pET vector containing a N or C-terminal 6xHis tag by Gibson assembly. The N-terminal intrinsically disordered intracellular region of GLPR (GLPR_IDR1_, residues R24-R214) was amplified by PCR using gDNA of *Cglu* as template and cloned into a pET vector containing an N-terminal 6xHis-SUMO tag by Gibson assembly. PCR products were digested with DpnI and transformed into chemio-competent *E. coli* cells. All plasmids were verified by Sanger sequencing (Eurofins).

### Cloning for recombinant protein expression in *C. glutamicum*

For ectopic recombinant expression of the different constructs in *Cglu*, we used the pUMS_3 shuttle expression vector ^7^, in which the gene of interest is placed under the control of *P_gntK_*, a tight promoter that is repressed by sucrose and induced by gluconate. Wild-type or mutant versions of the genes of interest were assembled in this plasmid by either Gibson assembly or site-directed mutagenesis using the primers listed in Table S6. For cellular localization studies, codon optimized mNeonGreen was ordered from Genscript and cloned alone or fused in frame to the N-terminus of GLP, MoeA1 (Cgl0212) and MoeA3 (Cgl1196) constructs including a GSGS flexible linker between the two fused proteins, or to the C-terminus of GLPR and Wag31 constructs. GLPR and GLPR_ΔIDR2_ expression vectors were generated from the pUMS3-GLPR-mNeon vector by introducing a STOP codon by PCR mutagenesis after GLPR residues 337 and 266 respectively.

### Protein expression and purification

N-terminal 6xHis-SUMO-tagged GLP and GLP-*Δ*Loop were expressed in *E. coli* BL21 (DE3) following an auto-induction protocol ^61^. After 4 hours at 37°C cells were grown for 20 hours at 20°C in 2YT supplemented with autoinduction medium and 50 µg/ml kanamycin. Cells were harvested and flash frozen in liquid nitrogen. Cell pellets were resuspended in 50 ml lysis buffer (PBS 1X, 500 mM NaCl, 20 mM Imidazole, benzonase, EDTA-free protease inhibitor cocktails (ROCHE)) at 4°C and sonicated. The lysate was centrifuged for 30min at 30000 x g at 4°C and loaded onto a Ni-NTA affinity chromatography column (HisTrap FF crude, GE Healthcare). His-tagged proteins were eluted with a linear gradient of buffer B (PBS 1X, 500 mM NaCl, 500 mM imidazole). The eluted fractions containing the protein of interest were pooled and dialyzed in presence or absence of the SUMO protease (ratio used 1:40). Dialysis was carried out at 18°C overnight in 20 mM Hepes pH 7.5, 500 mM NaCl. Cleaved his-tags and his-tagged SUMO protease were removed with Ni-NTA agarose resin. The SUMO-tagged or cleaved protein were concentrated and loaded onto a Superdex 200 16/60 size exclusion (SEC) column (GE Healthcare) pre-equilibrated at 4°C in 20 mM Hepes pH 7.5, 500 mM NaCl. The peak corresponding to the protein was concentrated, flash frozen in small aliquots in liquid nitrogen and stored at -80°C.

His-SUMO-FtsZ was purified as described ^7^. In brief, 6xHis-SUMO-FtsZ was produced as described above and cell pellets resuspended in 50 ml lysis buffer (50 mM Hepes pH 8, 300 mM KCl, 5% glycerol, 1 mM MgCl_2_, benzonase, lysozyme, 0.25 mM TCEP, EDTA-free protease inhibitor cocktails (ROCHE)) at 4°C and sonicated. The lysate was centrifuged for 30 min at 30000 x g at 4°C and loaded onto a Ni-NTA affinity chromatography column (HisTrap FF crude, GE Healthcare). His-tagged proteins were eluted with a linear gradient of buffer B (50 mM Hepes pH 8, 300 mM KCl, 5% glycerol, 1 M imidazole). The eluted fractions containing the protein of interest were pooled and dialyzed in presence or absence of the SUMO protease (ratio used 1:100). Dialysis was carried out at 4°C overnight in 25 mM Hepes pH 8, 150 mM KCl, 5% glycerol. Cleaved his-tags and his-tagged SUMO protease were removed with Ni-NTA agarose resin. The SUMO-tagged or cleaved protein was concentrated and loaded onto a Superdex 200 16/60 size exclusion (SEC) column (GE Healthcare) pre-equilibrated at 4°C in 25 mM Hepes pH 8, 150 mM KCl, 5% glycerol. The peak corresponding to the protein was concentrated, flash frozen in small aliquots in liquid nitrogen and stored at -80°C.

N- or C-terminal 6xHis-tagged GLPR was expressed in *E. coli* BL21 (DE3) using auto-induction. Cell pellets were resuspended in 150 ml lysis buffer (50 mM Hepes pH 7.5, 50 mM NaCl, 1 mM MgCl_2_, benzonase, EDTA-free protease inhibitor cocktails (ROCHE)) at 4°C and loaded 3 times in a CellD disrupter (Constant Systems). The lysate was centrifuged for 15min at 12000 x g at 4°C to remove cell debris and the supernatant was centrifuged again for 1h at 100000 x g at 4°C. The pellet containing the membrane fraction was resuspended in 40 ml membrane buffer (50 mM Hepes pH 7.5, 500 mM NaCl, 10 mM Imidazole, 10% glycerol, 1% DDM and EDTA-free protease inhibitor cocktails (ROCHE)) and incubated for 30min at 4°C. The membrane solubilized fraction was incubated for 1h at 4°C with 1 ml of Ni-NTA affinity chromatography beads (Super Ni-NTA resin, Neo Biotech). Beads were collected and washed with 10 column volumes of IMAC A buffer (50 mM Hepes pH 7.5, 300 mM NaCl, 25 mM Imidazole, 10 % glycerol, 0.05% DDM) and His-tagged GLPR was eluted with 10 column volumes of IMAC B buffer (50 mM Hepes pH 7.5, 300 mM NaCl, 500 mM Imidazole, 10 % glycerol, 0.05% DDM). The eluted fractions containing the protein of interest were pooled, concentrated, and loaded onto a Superdex 200 10/300 size exclusion (SEC) column (GE Healthcare) pre-equilibrated at 4°C in SEC buffer (25 mM Hepes pH 7.5, 150 mM NaCl, 10% glycerol and 0.05% DDM). The peak corresponding to GLPR was concentrated, flash frozen in small aliquots in liquid nitrogen and stored at -80°C.

6xHis-TEV-Wag31_1-61_ (DivIVA domain) was expressed in *E. coli* BL21 (DE3) using auto-induction. Cell pellets were resuspended in 50 ml lysis buffer (50 mM Hepes pH 7, 500 mM NaCl, 10 mM imidazole, 10% glycerol, benzonase, lysozyme, EDTA-free protease inhibitor cocktails (ROCHE)) at 4°C and sonicated. The lysate was centrifuged for 1h at 20000 x g at 4°C and loaded onto a Ni-NTA affinity chromatography column (HisTrap FF crude, GE Healthcare). His-tagged protein was eluted with a linear gradient of buffer B (50 mM Hepes pH 7, 500 mM NaCl, 10% glycerol, 1 M imidazole). The eluted fractions containing the protein of interest were pooled and dialyzed in the presence of the TEV protease (ratio 1:25). Dialysis was carried out at 18°C overnight in 50 mM Hepes pH 7, 150 mM NaCl, 5% glycerol. Cleaved his-tags and his-tagged TEV protease were removed with Ni-NTA agarose resin. The cleaved protein was concentrated and loaded onto a Superdex 200 16/60 size exclusion (SEC) column (GE Healthcare) pre-equilibrated at 4°C in 50 mM Hepes pH 7, 150 mM NaCl, 5% glycerol. The peak corresponding to the protein was concentrated, flash frozen in small aliquots in liquid nitrogen and stored at -80°C.

6xHis-SUMO-Wag31 was expressed in *E. coli* BL21 (DE3) using auto-induction. Cell pellets were resuspended in 50 ml lysis buffer (Hepes 20 mM pH 7, 150 mM NaCl, benzonase, EDTA-free protease inhibitor cocktails (ROCHE)) at 4°C and lysed by sonication. The lysate was centrifuged for 30min at 30000 x g at 4°C. After centrifugation, a gel-like layer containing His-SUMO-Wag31 was formed between the cell debris pellet and the clarified supernatant. This gel was recovered, washed 3 times with lysis buffer and solubilized in buffer Hepes 20 mM pH 8.5, NaCl 150 mM. Solubilized SUMO-Wag31 was digested overnight with SUMO protease (ratio used 1:100) at 18°C. Cleaved his-tags and his-tagged SUMO protease were removed with Ni-NTA agarose resin. Wag31 protein was dialyzed overnight at 4°C in buffer Hepes 20 mM pH 8.5, 150 mM NaCl and then concentrated, flash frozen in small aliquots in liquid nitrogen and stored at -80°C.

For antibody production N-terminal 6xHis-SUMO-GLPR_IDR1_ (corresponding to the intracellular IDR1 domain of GLPR) was expressed in *E. coli* BL21 (DE3) using auto-induction. Cell pellets were resuspended in 50 ml lysis buffer (Hepes 50 mM pH 8, 500 mM NaCl, 10 mM Imidazole, benzonase, EDTA-free protease inhibitor cocktails (ROCHE)) at 4°C and sonicated. The lysate was centrifuged for 30min at 30000 x g at 4°C and loaded onto a Ni-NTA affinity chromatography column (HisTrap FF crude, GE Healthcare). The His-tagged protein was eluted with a linear gradient of buffer B (Hepes 50 mM pH 8, 500 mM NaCl, 500 mM imidazole). The eluted fractions containing the protein of interest were pooled and dialyzed in the presence of the SUMO protease (ratio used 1:40). Dialysis was carried out at 18°C overnight in 20 mM Hepes pH 8, 150 mM NaCl. Cleaved his-tags and his-tagged SUMO protease were removed with Ni-NTA agarose resin. The cleaved protein was concentrated and loaded onto a Superdex 75 16/60 size exclusion (SEC) column (GE Healthcare) pre-equilibrated at 4°C in 20 mM Hepes pH 8, 150 mM NaCl. The peak corresponding to the protein was concentrated, flash frozen in small aliquots in liquid nitrogen and stored at -80°C. All purified proteins used in this work have been run on an SDS-PAGE and are represented in Figure S9.

### Crystallization

Crystallization screens were done using the sitting-drop vapor diffusion method and a Mosquito nanolitre-dispensing crystallization robot at 18°C (TTP Labtech, Melbourn, UK) following established protocols ^62^. Optimal crystals of GLP (13.5 mg/ml) were obtained in 100 mM Tris pH 8.5, 30% v/v PEG400 and 200 mM Na_3_Cit. For SAD phasing, GLP crystals were soaked in mother liquor containing 10 mM Cl_4_K_2_Pt for 30 min. The complex of GLP bound to the FtsZ_CTD_ peptide (DDLDVPSFLQ, purchased from Genosphere) was crystallized at 0.34 mM GLP (15 mg/ml) and 1.7 mM FtsZ_CTD_ (molar ratio 1:5). Crystals appeared after 2 weeks in 0.1M NaCl, 0.1M Bis-Tris pH 6.5 and 1.5M (NH4)_2_SO_4_. Crystals were cryo-protected in mother liquor containing 33% (v/v) ethylene glycol or 33% (v/v) glycerol.

### Data collection; structure determination and refinement

X-ray diffraction data were collected at 100 K using synchrotron radiation (Table S2) at Soleil (Gif-sur-Yvette, France) and processed using XDS ^63^ and AIMLESS from the CCP4 suite ^64^. A 2.35 Å dataset from a single crystal of GLP soaked in Cl_4_K_2_Pt was used to solve the crystal structure by SAD phasing using Phaser ^65^ and automatic model building with Buccaneer from the CCP4 suite. The structures of GLP alone and in complex with FtsZ_CTD_ were refined through iterative cycles of manual model building with COOT ^66^ and reciprocal space refinement with Phenix ^67^. The final crystallographic statistics and the PDB deposition codes of the atomic coordinates and structure factors are shown in Table S2. Structural figures were generated with ChimeraX ^68^.

### Differential Scanning Fluorescence (thermofluor) assay

For the thermofluor assay, 3 μg of GLP in 25 mM Hepes pH 8, 150 mM NaCl, 5% glycerol with or without 1 mM FtsZ_CTD_ were dispensed into a 96 well PCR plates (20 μl per well in triplicates). 0.6 μl a 50X Sypro Orange (Invitrogen) was added to each well and the mixture was heated from 25 to 95°C in 1°C steps of 1 min each in a CFX96 Touch™ Real-Time PCR Detection System (BioRad). Excitation and emission filters of 492 and 516 nm respectively, were used to monitor the fluorescence increase resulting from binding of the Sypro Orange to exposed hydrophobic regions of the unfolding protein. The midpoint of the protein unfolding transition was defined as the melting temperature T_m_.

### Bio-layer interferometry assays

Bio-layer interferometry experiments were performed on the Octet-Red384 device (Pall ForteBio) at 25°C. To test interactions between FtsZ and GLP variants, the His-SUMO_GLP variants were diluted at 227 nM in capture buffer (20 mM Hepes pH 7.5, 500 mM NaCl, BSA 1 mg/ml) and then immobilized on the commercially available Sartorius Ni–NTA biosensors for 10 minutes at 1000 rpm followed by a washing step of 2 min to remove any loosely bound protein. Empty sensors were used as reference for unspecific binding. FtsZ was diluted at 10 μM in polymerization buffer (25 mM Pipes pH 6.9, 100 mM KCl, 10 mM MgCl_2_) and pre-incubated with or without 1 mM GTP at room temperature for 20 min. Binding of monomeric or polymerized FtsZ filaments to the immobilized His-SUMO_GLP variants was monitored for 10 minutes with agitation at 1000 rpm followed by dissociation in the same buffer without proteins for 10 minutes.

In the reciprocal approach, His-SUMO-FtsZ was diluted at 4 μM in polymerization buffer supplemented with 1 mg/ml of BSA and then immobilized on the commercially available Sartorius Ni-NTA biosensors for 10 minutes at 1000 rpm shaking speed followed by a washing step in the same buffer of 3 min to remove any loosely bound protein. Sensors loaded with His-LysA (an unrelated protein from the mycobacteriophage TM4) was used as reference for non-specific binding. His-SUMO-FtsZ loaded, or reference sensors were incubated for 3 min at 1000 rpm in the absence and presence of two-fold serially diluted concentrations of GLP (80-1.25 μM range) in polymerization buffer followed by dissociation in the same buffer without protein for another 3 minutes.

To test the interaction between GLP, GLP-*Δ*Loop and GLPR, C-terminal His-tagged GLPR was immobilized at the surface of Ni-NTA biosensors and untagged GLP or GLP-*Δ*Loop were tested for binding to GLPR. Empty sensors were used as reference for unspecific binding. In these assays, GLPR was diluted at 0.4 μM in GLPR buffer (25 mM Hepes pH 7.5, 150 mM NaCl, 10% glycerol and 0.05% DDM, 1 mg/ml of BSA) and then immobilized on Ni-NTA biosensors for 10 minutes at 1000 rpm shaking speed followed by a washing step in the same buffer of 3 min to remove any loosely bound protein. Association of untagged GLP variants in GLPR buffer was monitored for 30 minutes followed by dissociation in the same buffer without proteins for another 30 minutes.

For the biotinylation reaction, 100 μl of His-GLPR at 25 μM was incubated with 20x molar excess of EZ-Link NHS-PEG4-Biotin (Thermo Scientific) following supplier instructions. Biotinylated GLPR was diluted to 0.25 μM in GLPR buffer (25 mM Hepes pH 7.5, 150 mM NaCl, 10% glycerol and 0.05% DDM) and then immobilized on the commercially available Sartorius Streptavidin biosensors for 5 minutes at 1000 rpm shaking speed followed by a washing step in GLPR buffer of 3 min to remove any loosely bound protein. Empty sensors were used as reference for non-specific binding.

GLPR loaded or empty reference sensors were incubated for 5 min at 1000 rpm in the absence and presence of two-fold serially diluted concentrations of Wag31 (150-2.34 μM range) in buffer A (20 mM Hepes pH 8.5, 150 mM NaCl, DDM 0.05%) or DivIVA (200-3.15 μM range) in buffer B (25 mM Hepes pH 7.5, 150 mM NaCl, 10% glycerol, 0.05% DDM, BSA 1 mg/ml). Specific signals were obtained by double referencing, that is, subtracting non-specific signals measured on reference sensors and buffer signals on specific loaded sensors.

Assays were performed at least twice, and to obtain the Kd value, steady-state signal versus concentration curves were fitted with GraphPad Prism 9 assuming a one site binding model.

### Circular Dichroism

All CD measurements were acquired with an Aviv 215 spectropolarimeter. Far-UV (195–260 nm) spectra were recorded at 25°C using a 0.2-mm path-length cylindrical cell. GLP variants were measured at 20 μM in a buffer containing 20 mM Hepes pH 7.5, NaCl 500 mM. Ellipticity was measured every 0.5 nm and averaged over 2 s. The final spectrum of the protein sample was obtained by averaging three successive scans and subtracting the baseline spectrum of the buffer recorded under the same conditions. Finally, BestSel ^69^ was used for quantitative decomposition of the far-UV CD spectrum.

Co-immunoprecipitation of Wag31/GLPR in *C. glutamicum*.

*Cglu*, *Cglu*_*Δglp* or *Cglu*_*Δglpr* strains were grown in 100 ml of CGXII minimal medium at 30°C for 6 hs. Cells were harvested, washed with 1X PBS and normalized by resuspending cell pellets in PBS-T (1X PBS, 0.1% v/v Tween-80) to give a final OD_600_ of 10. The cell suspensions were cross-linked with 0.25% v/v of formaldehyde for 20 min at 30°C with gentle agitation. The crosslinking reaction was stopped by adding 1.25 M glycine and incubated for 5 min at room temperature. Sample preparation and co-immunoprecipitation was performed as described above except that in this case magnetic agarose beads coupled with anti-GLPR antibodies were used. Eluted protein samples were subjected to immunoblot analysis using anti-GLPR or anti-DivIVA antibodies.

### Cell Fractionation

Analysis of protein subcellular localization was performed by cell disruption followed by differential centrifugation. *Cglu* strains were grown in 20 ml BHI medium at 30°C for 6 hs and harvested by centrifugation. To prepare cell extracts, bacterial cell pellets were resuspended in 1.5 ml of lysis buffer (25 mM Tris pH 7.5, 150 mM NaCl, benzonase, EDTA-free protease inhibitor cocktails (ROCHE)) and disrupted at 4°C with 0.1 mm glass beads and using a PRECELLYS 24 homogenizer. Cell debris and aggregated proteins were removed by centrifugation at 14000 × g for 20 min at 4°C. The supernatant was centrifuged again at 90000 × g for 30 min at 4°C to pellet cell membranes. Membrane fractions were solubilized with lysis buffer supplemented with 0.5% SDS. Protein concentrations were determined by UV_280_ absorbance and adjusted to 6 mg/ml. 120 μg of each fraction were run on an SDS-PAGE gel and analyzed by Western Blot using anti-GLP antibodies.

### Antibody production, purification and characterization

Anti-GLP, anti-GLPR, and anti-Wag31 antibodies were raised in rabbits and produced by Covalab using purified GLP, GLPR_IDR1_, or Wag31_1-61_ proteins as antigens. For antibody purification, rabbit serums from day 67 post-inoculation were purified using a 1 ml HiTrap NHS-Activated HP column (GE Healthcare) loaded with the corresponding antigen according to manufacturer instructions. 5 ml of rabbit serum were diluted in 1/3 in binding buffer (20 mM Sodium Phosphate pH 7.4, 500 mM NaCl) and loaded onto the antigen containing column, followed by a wash step with 7 ml of binding buffer. Antibodies were eluted with 10 ml of elution buffer (100 mM Glycine pH 3, 500 mM NaCl) and neutralized with 1M Tris pH 9. Purified antibodies were concentrated up to 8 mg/ml and then mixed 1:1 with glycerol 100%, aliquoted and stored at -20°C. Anti-SepF and anti-mScarlet antibody production was described previously ^7^.

### Western Blots

To prepare cell extracts, bacterial cell pellets were resuspended in lysis buffer (50 mM Bis-Tris pH 7.4; 75 mM 6-Aminocaproic Acid; 1 mM MgSO4; Benzonase and protease Inhibitor) and disrupted at 4°C with 0.1 mm glass beads and using a PRECELLYS 24 homogenizer. Total extracts (from 60 μg to 120 μg according to the protein studied) were run on a SDS-PAGE gel, electrotransferred onto a 0,2 μm nitrocellulose membrane and incubated for 1h with blocking buffer (5% skimmed milk in 1X TBS-Tween buffer) at room temperature (RT). Blocked membranes were incubated for 1 h at room temperature with the corresponding primary antibody diluted to the appropriate concentration in blocking buffer. After washing in TBS-Tween buffer, membranes were probed with an anti-rabbit or an anti-mouse horseradish peroxidase-linked secondary antibody (GE healthcare) for 45 minutes. For chemiluminescence detection, membranes were washed with 1X TBS-T and revealed with HRP substrate (Immobilon Forte, Millipore). Images were acquired using the ChemiDoc MP Imaging System (Biorad). All uncropped blots are shown in Figure S10.

### Mass spectrometry

#### Sample preparation for mass spectrometry analysis

For SepF interactome, *C. glutamicum* ATCC13032 (WT) and the strains expressing SepF-Scarlet, SepF_K125E/F131A_-Scarlet and Scarlet described previously ^7^ were used. All strains were grown in CGXII minimal media supplemented with 1% gluconate for 6 h at 30°C. Cells were harvested, washed, and normalized by resuspending cell pellets in PBS-T (1X PBS, 0.1% v/v Tween-80) to give a final OD_600_ of 3. The cell suspensions were cross-linked with 0.25% v/v of formaldehyde for 20 min at 30°C as previously described. Co-immunoprecipitation experiments (Co-IPs) were performed from protein extracts by incubation with magnetic beads cross-linked with 10 μg of purified antibodies (anti-Scarlet or anti-SepF produced by Covalab) following the protocols previously described ^7^. Briefly, proteins recovered after washing an elution with 1 M glycine pH 2 were neutralized with 1 M Tris pH 9, denatured (2M urea), reduced (10 mM DTT, 1h, RT), alkylated (55 mM IAM, 45 min, RT, in the dark) and digested with 0.5 μg of trypsin (Promega). Tryptic peptides were desalted using a POROS^TM^ R2 resin (ThermoFisher), vacuum dried and resuspended in 0.1% formic acid (FA). Four replicates for each condition were analysed by nano-HPLC-MS/MS.

GLP interactomes were analysed in two strains: *Cglu* and *Cglu*_*Δglpr*, using the *Cglu*_*Δglp* as a control. Strain growth and cross-linking were performed as described above. Co-immunoprecipitation experiments were performed using anti-GLP antibodies. Eluted proteins were neutralized (1 M Tris pH 9), denatured (6M urea), reduced and alkylated with IAM 55 mM before digestion with 0.5 μg of Trypsin (Promega) using the FASP protocol ^70^ with filter passivation in 5% Tween-20 ^71^ was used for sample preparation. Tryptic digests were desalted using ZipTips C18 (Merck Millipore), vacuum dried and resuspended in 0.1% formic acid. Tryptic peptides from 4 replicates of each condition were analyzed using a nano-HPLC-MS/MS.

To calculate protein enrichment, we analysed the full proteome of *Cglu* grown in CGXII minimal media. Protein extracts were run on 1 cm long SDS-PAGE gels (12% acrylamide). In-gel Cys alkylation was performed by incubation with DTT and IAM as described before. After in-gel digestion, peptide mixtures were desalted using ZipTips C18 (Merck Millipore), vacuum dried and resuspended in 0.1% formic acid. 3 replicates were analysed using a nano-HPLC-MS/MS.

#### Nano-HPLC-MS/MS

Tryptic peptides were analysed using a nano-HPLC (UltiMate 3000, Thermo) coupled to a hybrid quadrupole-orbitrap mass spectrometer (QExactive Plus, Thermo). Peptide mixtures were loaded on C18 columns and separated using a two-solvent system: (A) 0.1% FA in water and (B) 0.1% FA in acetonitrile (ACN) at a flow rate of 200 nL/min. Capillary temperature was set at 250°C and spray voltage ranged from 1.7 to 2 kV. The survey scans were acquired in a range of 200-2000 m/z with a resolution of 70000 at 200 m/z, an AGC target value of 1E6 and a maximum ion injection time of 100 ms. Precursor fragmentation occurred in an HCD cell with a resolution of 17500 at 200 m/z, an AGC target value of 1E5 and a maximum ion injection time of 50 ms. Normalized collision energy was used in steps of NCE 25, 30 and 35. Online MS analysis was carried out in a data-dependent mode (MS followed by MS/MS of the top 12 ions) using dynamic exclusion. Chromatography conditions for SepF interactome were described previously ^7^. Briefly, peptide mixtures were separated on a C18 column (PepMap® RSLC, 0.075 × 500 mm, 2 μm, 100 Å) using a 65 min gradient of mobile phase B from 0 to 55%. For GLP co-IPs, peptide mixtures were loaded onto a pre-column (Acclaim PepMapTM 100, C18, 75 µm X 2 cm, 3 µm particle size) and separated with an Easy-Spray analytical column (PepMapTM RSLC, C18, 75 µm X 50 cm, 2 µm particle size) using an elution gradient from 1% to 35% B over 90 min followed by a 35% to 99% B step over 20 min. For total proteome analysis, elution was achieved using a separation gradient of 150 min from 1% to 35%B.

#### Protein identification and data analysis

PatternLab for Proteomics V software (PatternLabV) was used to perform peptide spectrum matching and label-free quantitation analyses based on extracted-ion chromatogram (XIC) ^72^. For the SepF interactome analysis a target reverse database was generated by PatternLab using the *Cglu* ATCC13032 proteome downloaded from UniProt (November 2018) to which the sequences of Scarlet, SepF-Scarlet, SepF_K125E/F131A_-Scarlet and the most common contaminants in proteomics experiments were added. For the GLP interactome, a target reverse database obtained from *Cglu* ATCC13032 proteome downloaded from UniProt (November 2021) and the most common contaminants in proteomics was used. *Cglu* ATCC13032 database was also use for proteome analysis.

Search parameters were set as follows: m/z precursor tolerance: 35 ppm, methionine oxidation and cysteine carbamidomethylation as variable and fixed modifications respectively, and a maximum of 2 missed cleavages and 2 variable modifications per peptide. Search results were filtered by the PatternLab Search Engine Processor (SEPro) algorithm with a maximum FDR value ≤ 1% at protein level and 10 ppm tolerance for precursor ions.

To identify SepF interactors, we compared the list of proteins recovered under different conditions: WT strain using *α*-SepF antibodies (WT/*α*-SepF), SepF-Scarlet strain using *α*-SepF antibodies (SepF-Scarlet/*α*-SepF) and SepF-Scarlet strain using *α*-Scarlet antibodies (SepF-Scarlet/*α*-Scarlet). As a control of background binding, the Scarlet strain using *α*-Scarlet antibodies was used. Additionally, we evaluated the recovery of the core SepF interactors using a SepF mutant (SepF_K125E/F131_), with impaired binding for FtsZ. For that purpose, we compared proteins recovered from SepF-Scarlet and SepF_K125E/F131A_-Scarlet using *α*-Scarlet antibodies.

In a similar way, to identify GLP interactors we compared the proteins recovered from *Cglu* or *Cglu*_*Δglpr* with *Cglu*_*Δglp* strain, using four replicates of each strain.

To compare proteins identified in co-IPs with control, PatternLab’s Venn diagram statistical module was used. This module allows to determine proteins uniquely detected in each biological condition using a probability value <0.05 ^73^. PatternLab V was also used to relatively quantify proteins using XIC. Pairwise comparison between proteins recovery from Co-IPs and controls was performed using the XIC browser mode and the following conditions: maximum parsimony, minimum number of peptides: 1, minimum number of MS1 counts: 5, considering only preferred charge state and Log2FC > 1.8. This module analyses differences at the peptide level and uses the Benjamini-Hochberg’s theoretical estimator to deal with multiple T-tests. Enrichment factors for SepF interactors were calculated as the ratio of the NSAF of each interactor in the interactome and in the proteome. To compare the recovery of SepF interactors in pull down analyses of SepF-Scarlet and SepF_K125E/F131A_-Scarlet strains, the signal of each interactor (ΣXIC signal of detected peptides in each replicate) was normalized by the signal of SepF in the corresponding sample. The statistical analysis was performed using unpaired Student’s t-test (p < 0.05). All data is presented as mean ± SD. Calculations were done using GraphPad Prism. The same approach was used to compare the recovery of Wag31 in GLP interactome of *Cglu* and *Cglu*_*Δglpr* strains.

#### Phase contrast and fluorescence microscopy

For imaging, cultures of *C. glutamicum* were grown in BHI for around 6 hours, then pelleted at 5200 x g at room temperature, washed with 0.9% NaCl and inoculated into CGXII media supplemented with 4% sucrose and kanamycin (25 μg/mL) for overnight growth. The following day cultures were diluted to OD_600_ of 1 in CGXII with 4% sucrose (with or without 1% gluconate) and grown for 6 hours to a required OD_600_ of about 4-6 (early exponential phase). For each sample, 100 μL of culture were pelleted, washed with fresh medium and diluted to an OD_600_ of 3 for imaging. For membrane staining, Nile Red (Enzo Life Sciences) was added to the culture (2 μg/ml final concentration) just prior to placing them on 2% agarose pads prepared with the corresponding growth medium. Cells were visualized using a Zeiss Axio Observer Z1 microscope fitted with an Orca Flash 4 V2 sCMOS camera (Hamamatsu) and a Pln-Apo 63X/1.4 oil Ph3 objective. Images were collected using Zen Blue 2.6 (Zeiss) and analyzed using the software Fiji ^74^ and the plugin MicrobeJ ^75^ to generate violin plots and fluorescent intensity heat maps. For all analyses, the *Cglu* strain corresponds to *Cglu* + empty plasmid.

Because of the important number of cells analyzed in each sample, Cohen’s *d* value was used to describe effect sizes between different strains independently of sample size:

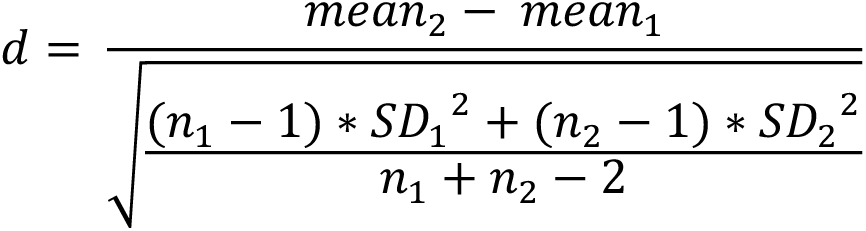

Values were interpreted according to the intervals of reference suggested by Cohen ^76^ and expanded by Sawilowsky ^77^, as follows: small (), *d* < 0.50; medium (*), 0.50 < *d* < 0.80; large (**), 0.80 < *d* <1.20; very large (***), 1.20 < *d* < 2.0; huge (****), *d* > 2.0.

*p* values were obtained by a student test calculated on R (Welch two sample t-test). The experiments were performed as biological triplicates. Some autofluorescence is observed for wild-*type Cglu* as previously described ^7^.

#### Protein database assemblies

To carry out the sequence analyses in Actinobacteria, we assembled two databases representing all Actinobacteria diversity present at the National Center for Biotechnology (NCBI) as of February 2021: ACTINO_DB and ACTINO_REDUCED_DB. ACTINO_DB contains 244 taxa; we selected five species per class, except for class Corynebacteriales, where we selected five species per order (for a list of taxa see Table S3). ACTINO_REDUCED_DB contains 113 taxa from ACTINO_DB (for a list of taxa see Table S3). For the phylogenetic analyses in Bacteria, we assembled a database based on the one provided in ^78^. We reduced the taxonomic sampling to 76 species by removing all candidate phyla (for a list of taxa see Table S4). Finally, for the phylogenetic analyses in Archaea, we worked with the same database provided in ^79^, consisting of 155 species, representatives of all archaeal diversity.

#### Homology searches and mapping

To identify all MoeA homologs in the ACTINO_DB we used HMM profile searches. First, we used the HMMER package (v3.3.2) ^80^ tool jackhmmer to look for homologs of GLP and GLPR in all the proteomes using the GenBank ^81^ sequences BAB98276.1 and BAB98278.1 as query respectively. The hits were aligned with mafft (v7.475) ^82^ using default parameters. The alignments were manually curated, removing sequences that did not align. The hits obtained by jackhmmer might not include sequences that are very divergent from the single sequence query. For this reason, the curated alignments were used to create HMM profiles using the HMMER package tool hmmbuild. These curated HMM profiles for GLP and GLPR were used for a final round of searches against the ACTINO_DB, Bacteria and Archaea databases, using the HMMER tool hmmsearch. All hits for each search were curated to remove false positives by checking alignments obtained using linsi, the accurate option of mafft (v7.475). For Actinobacteria, hits were also curated based on their genomic context. First, we retrieved five genes upstream and downstream each MoeA paralog identified in Actinobacteria. Then, we grouped the corresponding proteins into protein families. Each family larger than 10 sequences was used to create HMM profiles as explained before. These profiles together with the GLP and GLPR profiles were used in MacSyFinder ^83^ against the ACTINO_DB to identify conserved genomic contexts containing MoeA and three or more members of these families separated by no more than five other proteins and a permissive evalue (< 0.1). This analysis also complemented the GLPR homology searches, as the sequences are very divergent and therefore difficult to identify if it is not because of their genomic context. Finally, we analyzed the taxonomic distribution of the identified GLP and GLPR sequences. The phyletic pattern and the genomic context information was mapped on an Actinobacteria reference phylogeny using the online tool iTOL ^84^ and custom scripts.

#### Phylogenetic analyses

The alignments of GLP homologs (that includes all MoeA paralogs) were trimmed using BMGE (v1.2) ^85^ (option -m BLOSUM30) to keep only informative positions. These alignments were used to reconstruct the phylogeny of the MoeA paralogs in ACTINO_REDUCED_DB, Bacteria and Archaea. We used the maximum-likelihood phylogeny reconstruction tool IQ-TREE (v2.0.6) ^86^, with the LG+F+R8, LG+F+R10 and LG+R8 models respectively (-m MFP) and ultrafast bootstraps (-B 1000).

To reconstruct the reference phylogeny of ACTINO_DB we concatenated proteins RNApol subunits B and B’, and IF-2. Homologs of these proteins were identified, aligned and trimmed as explained before. These alignments were concatenated into a supermatrix that was used to infer a maximum likelihood (ML) tree with IQ-TREE, using the posterior mean site frequency (PMSF), and the model LG+C60+F+I+G with ultrafast bootstrap supports calculated from 1,000 replicates. The guide tree required by the PMSF model was previously obtained using the MFP option and the same supermatrix. The reference phylogeny of Bacteria was obtained from ^78^, and candidate taxa were pruned from the tree. The reference phylogeny of Archaea was obtained from ^79^.

## Acknowledgements

We gratefully acknowledge the core facilities at the Institut Pasteur C2RT, P. England, B. Raynal, S. Brûlé (PFBMI); P. Weber, C. Pissis, A. Mechaly (PFC), and J. Fernandes and A. Salles (UtechS PBI / Imagopole, supported by France BioImaging; ANR-10–INSB–04; Investments for the Future). We thank P. Campagne from the Bioinformatics and Biostatistics Hub from the IP-C3BI. We also acknowledge the synchrotron source Soleil (Saint-Aubin, France) for granting access to the facility and the staff of Proxima 1 and Proxima 2A beamlines for helpful assistance during X-ray data collection. Molecular graphics were done with ChimeraX, developed at UCSF with support from NIH (R01-GM129325) and NIAID. This work was supported in part by grants from the Agence Nationale de la Recherche (ANR, France), contracts ANR-18-CE11-0017 (P.M.A.) and ANR-21-CE11-0003 (A.M.W.), Agencia Nacional de Investigacion et Innovacion (ANII, Uruguay), FCE_1_2019_1_155569 (R.D.), FOCEM - Fondo para la Convergencia Estructural del Mercosur, COF 03/11 (R.D.), ECOS-Sud France-Uruguay, contract U20B02 (A.M.W. and R.D.), and by institutional grants from the Institut Pasteur, the CNRS, and Université Paris Cité. J.P. was funded through the AMX program from the Ecole Polytechnique. A.S. was part of the Pasteur - Paris University (PPU) International PhD Program, funded by the European Union’s Horizon 2020 research and innovation programme under the Marie Sklodowska-Curie grant agreement No 665807. Q.G. was funded by MTCI PhD school (ED 563). A.R was funded by ANII.

## Author contributions

A.M.W., R.D and P.M.A. designed the research. M.M., J.P., A.L., A.S. Q.G., M.B.A. and A.M.W. conducted the protein biochemistry, cell biology and genetic experiments, and purified proteins for structural and biophysical studies. J.P., A.D., C.G. and A.M.W. performed cellular imaging and analysis. M.M. and J.P. carried out the biochemical and biophysical studies of protein-protein interactions. A.L., M.M., A.R., M.P., A.M.W, and R.D. carried out MS and interactomics experiments. M.M., A.S. A.H., and P.M.A. carried out the crystallogenesis and crystallographic studies. D.M. performed the phylogeny and sequence analyses. A.M.W. and P.M.A. wrote the paper. All authors edited the paper.

## Data availability

Atomic coordinates and structure factors have been deposited in the PDB with accession codes 8BVE (GLP) and 8BVF (GLP-FtsZ_CTD_). The mass spectrometry proteomics data have been deposited to the ProteomeXchange Consortium via the PRIDE ^87^ partner repository with the dataset identifier PXD037255. All phylogenetic data used to produce our results is provided as Supporting Data and can be found under the following link: https://data.mendeley.com/datasets/265wyk8r3f/draft?a=6d0ecd8f-0adb-4f71-b984-37b7710c3f0a.

## FIGURE LEGENDS

