## Supplementary Information for "Eukaryotic-like gephyrin and cognate membrane receptor coordinate corynebacterial cell division and polar elongation"

**This PDF file includes:**

Figures S1 to S10

Tables S2, S5 and S6

Supplementary References

Note: Tables S1, S3 andS4 are provided separately as excel file

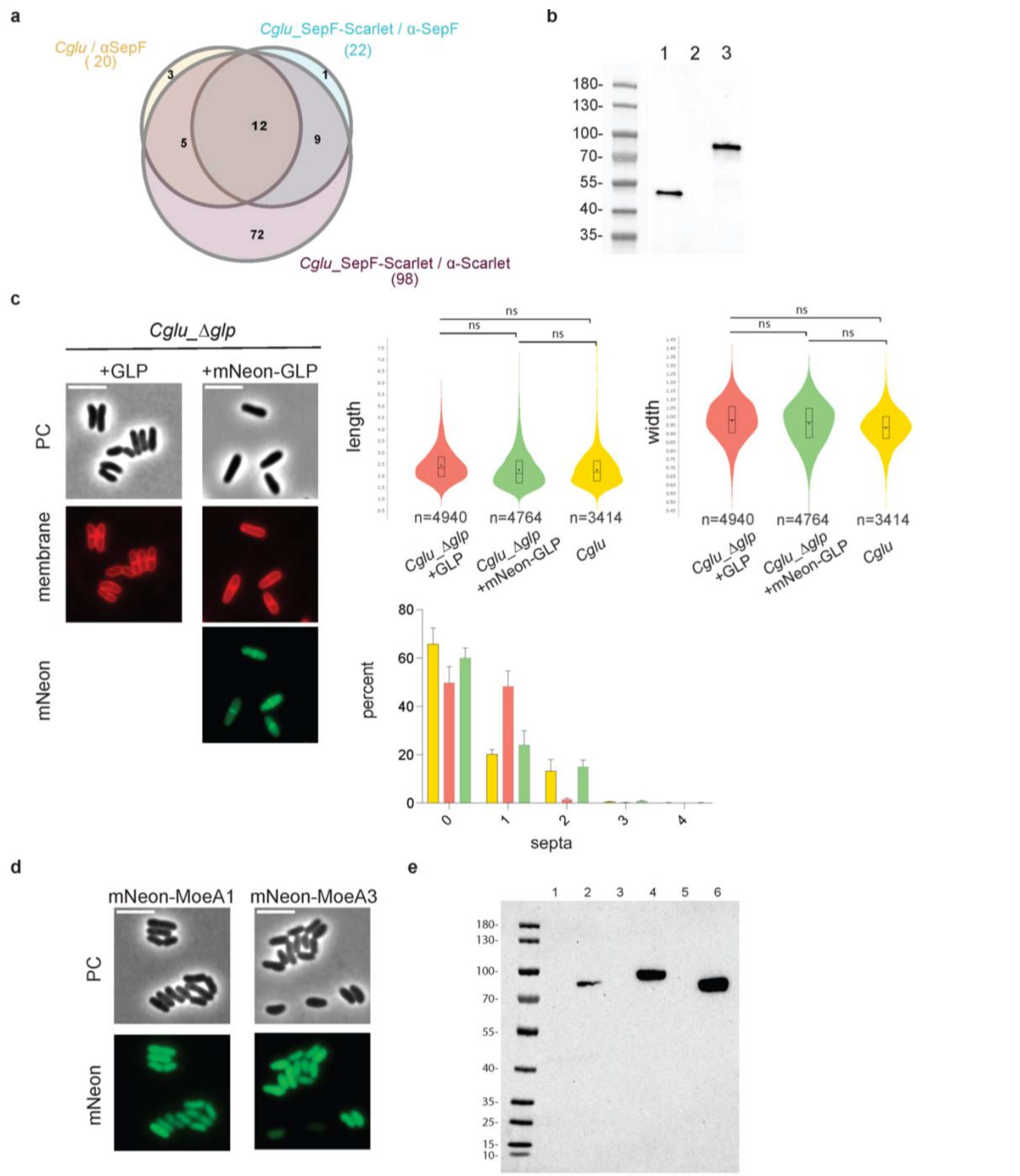

**Figure S1**. **Identification of GLP as a cell division protein.** (**a**) Venn diagram showing the overlap between three independent SepF interactomes using *Cglu* or *Cglu*_SepF-Scarlet strains. Proteins only detected in each interactome were identified by comparison with control condition and using the probability mode of Patternlab Venn diagram module (*p* value <0.05). Proteins enriched in SepF Co-IPs when compared to controls were identified using pairwise comparison module of Patternlab V based on XIC intensities. Twenty, 22 and 98 proteins were detected as SepF interactors in *Cglu*/α-SepF (strain/antibody), *Cglu*_SepF-Scalet/α-SepF and *Cglu*_SepF-Scalet/α-Scarlet respectively. Twelve proteins were common to all of Co-IPs, and for eleven of them an enrichment factor in relation to the total proteome could be calculated, and thus represent the core SepF interactome (Table S1a and Figure 1a). One additional interactor, the hypothetical protein Cgl1805, could not be detected in the proteome and no enrichment factor could thus be reported. (**b**) Western blots of whole cell extracts (120 μg) from *Cglu* (lane 1) and *Cglu_Δglp* (lane 2) strains complemented with the empty plasmid or mNeon-GLP (lane 3). GLP and mNeon-GLP levels were revealed using an α-GLP antibody. (**c**) Complementation of *Cglu_Δglp*. Left, representative images in phase contrast, Nile Red (membrane) and mNeon fluorescent signal for the *Cglu_Δglp* strain complemented with GLP or mNeon-GLP. Top right, violin plots showing the distribution of cell length and width for *Cglu_Δglp* + GLP (red), *Cglu_Δglp* + *Cglu*_mNeon-GLP (green) and *Cglu* (yellow). Significance indicated corresponds to values of Cohen's d (for length, from top to bottom: (ns, d = 0,29, p = 3,78e-37), (ns, d = 0,27, p = 2,59e-39), (ns, d = 0, p = 0,95); for width, from top to bottom: (ns, d = 0,38, p = 9,65e-74), (ns, d = 0,08, p = 1,74e-12), (ns, d = 0,25, p = 4,13e-25)). Frequency histogram indicating the number of septa per cell for *Cglu* (yellow), *Cglu_Δglp* + GLP (red) and *Cglu_Δglp* + mNeon-GLP (green) strains, calculated from n cells imaged from 3 independent experiments (triplicates) for each strain (for *Cglu*, n=718, 1468 and 1223; for *Cglu_Δglp* + GLP*,* n=2465, 1169 and 1297; for *Cglu_Δglp* + mNeon-GLP, n=1641, 1311 and 1801); bars represent the mean ± SD. (**d**) Cellular localization of *Cglu* MoeA homologs. Representative images in phase contrast and mNeon-MoeA1 / mNeon-MoeA3 fluorescent signal for *Cglu*_mNeon-MoeA1 and *Cglu*_mNeon-MoeA3. Both MoeA1 and MoeA3 are cytosolic, which contrasts with the mid-cell localization of mNeon-GLP shown in Figure 1d. All Scale bars 5μm. For all violin plots: The box indicates the 25$^{th}$ to the 75$^{th}$ percentile, the mean and the median are indicated with a dot and a line in the box, respectively. The number of cells used in the analyses (n) is indicated below each violin representation and correspond to triplicates.

UFB > 0.85. The scale bar represents the average number of substitutions per site. Branches that correspond to *C. glutamicum* and *M. tuberculosis* species are indicated in red.

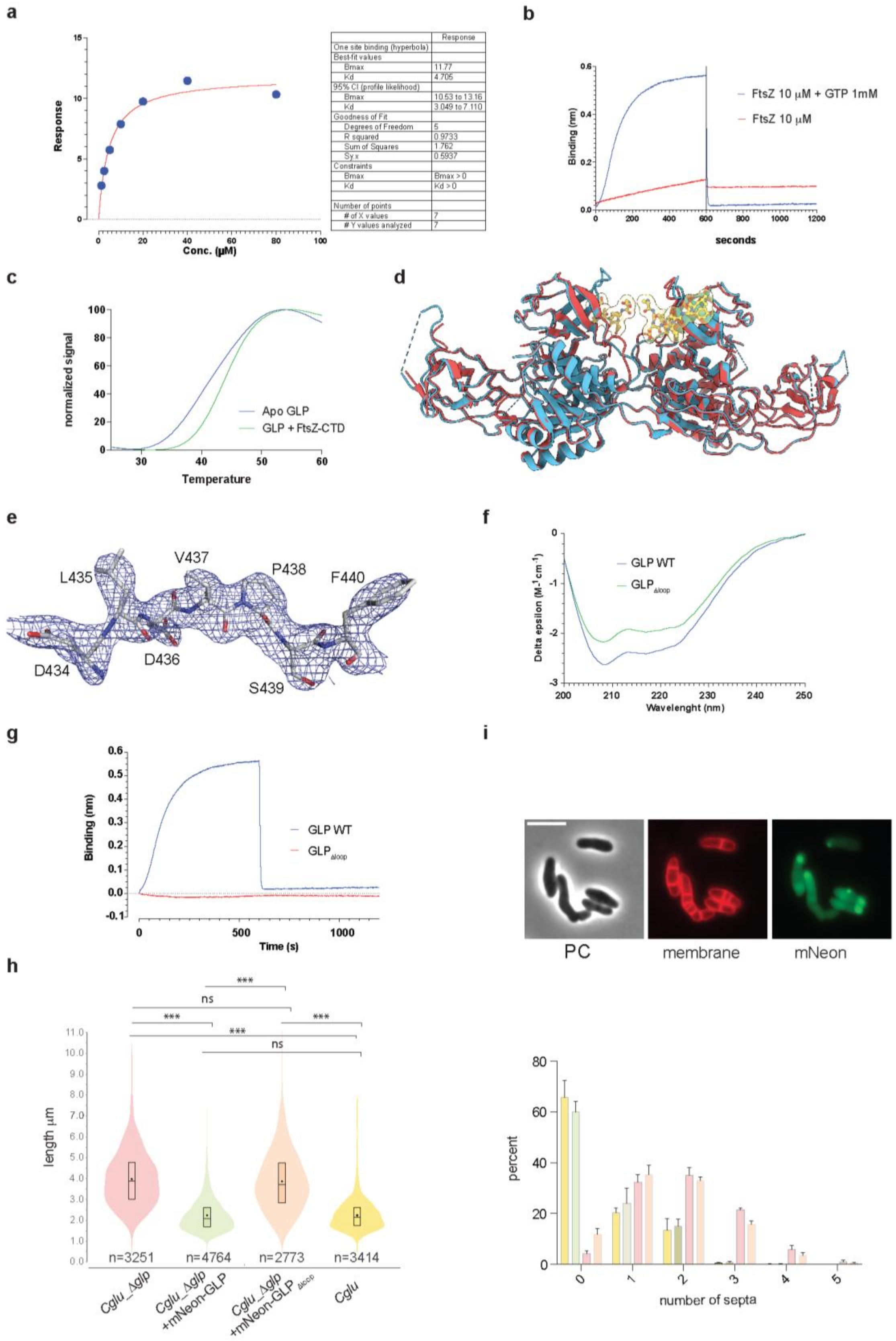

**Figure S3. GLP-FtsZ interaction.** (**a**) BLI data analysis for the GLP-FtsZ association profiles shown in Figure 2b. To obtain the *Kd* value (4.7 µM), steady-state signal versus concentration curves were fitted assuming a one-site binding model. (**b**) BLI sensorgrams of FtsZ binding to immobilized GLP in the presence or absence of 1 mM GTP. (**c**) Normalized melting curves of GLP with or without 1 mM $FtsZ_{CTD}$ peptide as determined by a thermofluor assay. GLP was stabilized by 2 °C in the presence of $FtsZ_{CTD}$. (**d**) The crystal structures of ligand-free (red) and $FtsZ_{CTD}$-bound (blue) GLP can be superimposed with an r.m.s.d. of 0.64 Å for 392 equivalent $C\alpha$ atoms. (**e**) Electron density map of $FtsZ_{CTD}$ bound to GLP monomer B contoured at 1.2 $\sigma$. (**f**) Far-UV circular dichroism spectra of GLP and $GLP_{\Delta loop}$. (**g**) BLI sensorgrams of FtsZ binding to immobilized GLP or $GLP_{\Delta loop}$. (**h**) Complementation with $GLP_{\Delta loop}$. Left panel, violin plots showing the distribution of cell length for *Cglu_*Δ*glp* (red), *Cglu_*Δ*glp* + mNeon-GLP (green), *Cglu_*Δ*glp* + mNeon-$GLP_{\Delta loop}$ (orange) and *Cglu* (yellow). The number of cells used in the analyses (n) is indicated below each plot representing triplicate experiments. The box indicates the 25$^{th}$ to the 75$^{th}$ percentile, the mean and the median are indicated with a dot and a line in the box, respectively. Significance indicated corresponds to values of Cohen's d (from top to bottom: (***, d = 1,58, p = 0), (ns, d = 0,09, p = 0,0012), (*** left, d = 1,78, p = 0), (*** right, d = 1,54, p = 0), (***, d = 1,76, p = 0), (ns, d = 0, p = 0,95)). Right panel, frequency histogram indicating the number of septa per cell for *Cglu* (yellow), *Cglu_*Δ*glp* + mNeon-GLP (green), *Cglu_*Δ*glp* (red) and *Cglu_*Δ*glp* + mNeon-$GLP_{\Delta loop}$ (orange) strains, calculated from n cells imaged from 3 independent experiments (triplicates) for each strain (for *Cglu*, n=718, 1468 and 1223; for *Cglu_*Δ*glp* + mNeon-GLP, n=1641, 1311 and 1801; for *Cglu_*Δ*glp*, n=873, 1538 and 840; for *Cglu_*Δ*glp* + mNeon-$GLP_{\Delta loop}$ , n=678, 1187 and 905); bars represent the mean $\pm$ SD. (**i**) Cellular localization of *Cglu_*Δ*glp* + mNeon-$GLP_{\Delta loop}$. Representative images for phase contrast, mNeon-$GLP_{\Delta loop}$ and nile red (membrane). Scale bar = 5$\mu$m.

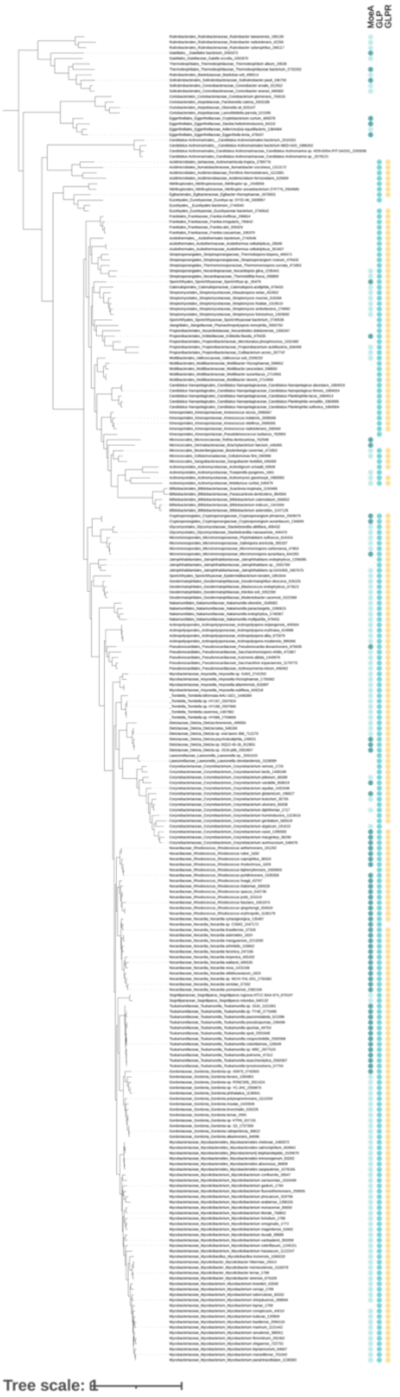

**Figure S4. Phyletic pattern for the presence of MoeA, GLP and GLPR in Actinobacteria.** Extended version of Figure 3. Full circles indicate presence of the gene in the species and blanks indicate its absence. In the column MoeA, darker blue indicates the presence of more than one copy. Column MoeA represents all paralogs, except for GLP that is indicated in a separate column. The phyletic pattern is represented on a reference Actinobacteria tree. Dots indicate UFB > 0.85. The scale bar represents the average number of substitutions per site.

a

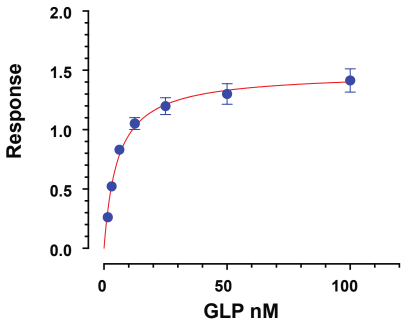

| | |
|---|---|
| One site binding (hyperbola) | |
| Best-fit values | |
| Bmax | 1.479 |
| Kd | 5.547 |
| 95% CI (profile likelihood) | |
| Bmax | 1.405 to 1.557 |
| Kd | 4.549 to 6.741 |
| Goodness of Fit | |
| Degrees of Freedom | 12 |
| R squared | 0.9805 |
| Sum of Squares | 0.04260 |
| Sy.x | 0.05958 |
| Constraints | |
| Bmax | Bmax > 0 |
| Kd | Kd > 0 |

b

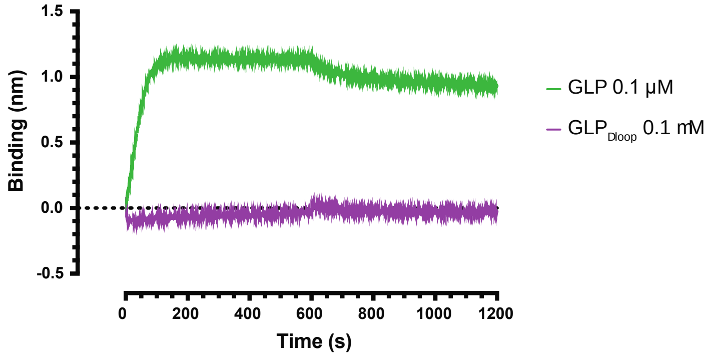

**Figure S5. GLP-GLPR interactions.** (**a**) BLI data analysis for the GLP-GLPR complex. To obtain the *Kd* value (5.5 nM) from the interaction profiles shown in Figure 4b, steady-state signal versus concentration curves were fitted assuming a one site binding model. (**b**) BLI sensorgrams of GLP or $GLP_{\Delta loop}$ (0.1 µM) binding to immobilized GLPR.

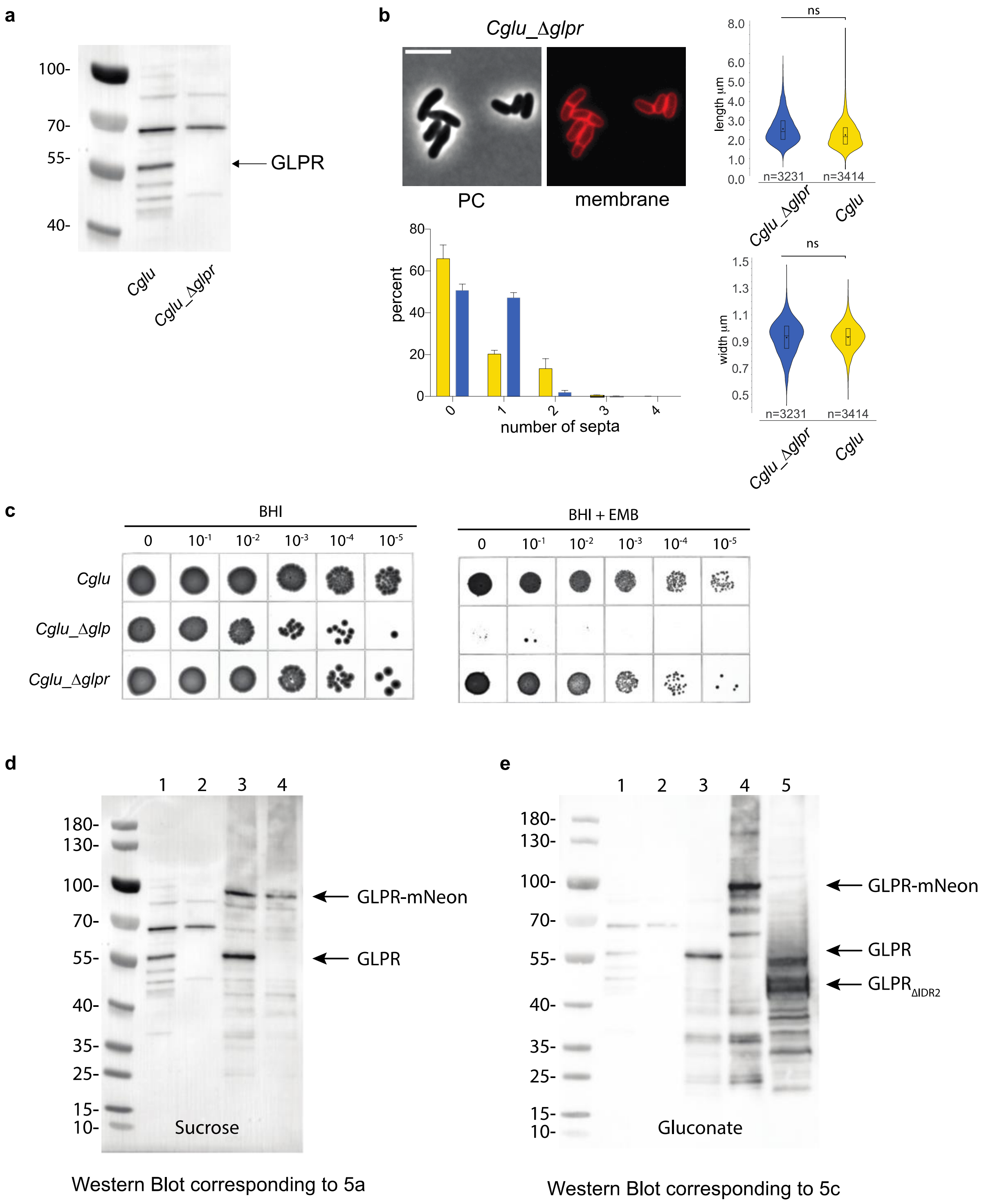

**Figure S6. Phenotypic analysis of the *Cglu_Δglpr* strain.** (**a**) Western blots of whole cell extracts (120 µg) from wild-type *Cglu* and *Cglu_Δglpr*. GLPR levels were revealed using an α-GLPR antibody. An arrow indicates the specific signal for GLPR in the blot. (**b**) Representative images in phase contrast and membrane staining by Nile Red fluorescent signal of the *Cglu_Δglpr* . Scale bars 5µm. Violin plots showing the distribution of cell length (ns, d = 0,46, p = 8,22e-75) and cell width (ns, d = 0, p = 0,29) for *Cglu_Δglpr*  (blue) and *Cglu* (yellow) at time point 6 hours in minimal media; the number of cells used (from triplicates) in the analyses (n) is indicated below each violin representation; the box indicates the $25^{th}$ to the $75^{th}$ percentile, the mean and the median are indicated with a dot and a line in the box, respectively. Frequency histogram indicating the number of septa per cell for *Cglu* (yellow) and *Cglu_Δglpr* (blue) strains, calculated from n cells imaged from 3 independent experiments for each strain (for *Cglu*, n=718, 1468 and 1223; for *Cglu_Δglpr,* n=931, 934 and 1363); bars represent the mean ± SD. (**c**) Ethambutol sensitivity assay of Δ*glpr strain*. BHI overnight cultures of *Cglu*, Δ*glp and* Δ*glpr* strains were normalized to an $OD_{600}$ of 0.5, serially diluted 10-fold, and spotted (10 µl) onto BHI agar medium with and without 1 µg/ml EMB as indicated. Plates were incubated for 48 hours at 30°C and photographed. (**d**) Western blots of whole cell extracts (120 µg) from *Cglu* and *Cglu_Δglpr* strains complemented with the empty plasmid or GLPR-mNeon. GLPR levels were revealed using an α-GLPR antibody. An arrow indicates the specific signal for GLPR and GLPR-mNeon in the blot. Western blot corresponds to the representative cells shown in Figure 5a. Lane 1: *Cglu* + empty plasmid; Lane 2: *Cglu_Δglpr* + empty plasmid; Lane 3: *Cglu* + GLPR-mNeon; Lane 4: *Cglu_Δglpr* + GLPR-mNeon. (**e**) Western blots of whole cell extracts (120 µg) from *Cglu* and *Cglu_Δglpr* strains complemented with the empty plasmid, GLPR , GLPR-mNeon or $GLPR_{ΔIDR2}$. GLPR levels were revealed using an α-GLPR antibody. Arrows indicate specific signal for GLPR, GLPR-mNeon and $GLPR_{ΔIDR2}$ in the blot. Western blot corresponds to the representative cells shown in Figure 5c. Lane 1: *Cglu* + empty plasmid; Lane 2: *Cglu_Δglpr* + empty plasmid; Lane 3: *Cglu_Δglpr* + GLPR; Lane 4: *Cglu_Δglpr* + GLPR-mNeon; Lane 5: *Cglu_Δglpr* + $GLPR_{ΔIDR2}$.

**a**

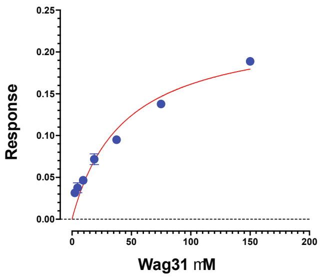

| | |
|---|---|
| One site binding (hyperbola) | |
| Best-fit values | |
| Bmax | 0.2314 |
| Kd | 43.38 |
| 95% CI (profile likelihood) | |
| Bmax | 0.1924 to 0.2882 |
| Kd | 26.54 to 72.70 |
| Goodness of Fit | |
| Degrees of Freedom | 12 |
| R squared | 0.9509 |
| Sum of Squares | 0.002011 |
| Sy.x | 0.01295 |
| Constraints | |
| Bmax | Bmax > 0 |
| Kd | Kd > 0 |
| Number of points | |
| # of X values | 14 |
| # Y values analyzed | 14 |

**b**

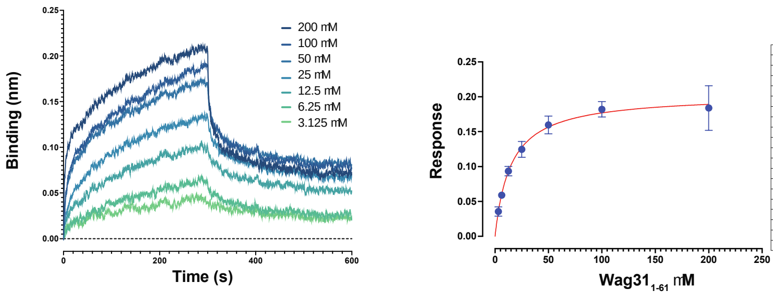

| | |
|---|---|
| One site binding (hyperbola) | |
| Best-fit values | |
| Bmax | 0.2032 |
| Kd | 14.86 |
| 95% CI (profile likelihood) | |
| Bmax | 0.1869 to 0.22 11 |
| Kd | 10.96 to 20.01 |
| Goodness of Fit | |
| Degrees of Freedom | 12 |
| R squared | 0.9613 |
| Sum of Squares | 0.001694 |
| Sy.x | 0.01188 |
| Constraints | |
| Bmax | Bmax > 0 |
| Kd | Kd > 0 |
| Number of points | |
| # of X values | 14 |
| # Y values analyzed | 14 |

**Figure S7. GLPR-Wag31 interactions.** (**a**) BLI data analysis for the GLPR-Wag31 association. To obtain the *Kd* value (43.3 µM) from the interaction profiles shown in Figure 5j, steady-state signal versus concentration curves were fitted assuming a one site binding model. (**b**) Sensorgrams of $Wag31_{1-61}$ binding to immobilized GLPR by biolayer interferometry. A series of measurements using a range of concentrations for $Wag31_{1-61}$ was carried out to derive the apparent equilibrium dissociation constant *Kd* (14.86 µM) from steady-state signal versus concentration curves fitted assuming a one-site binding model.

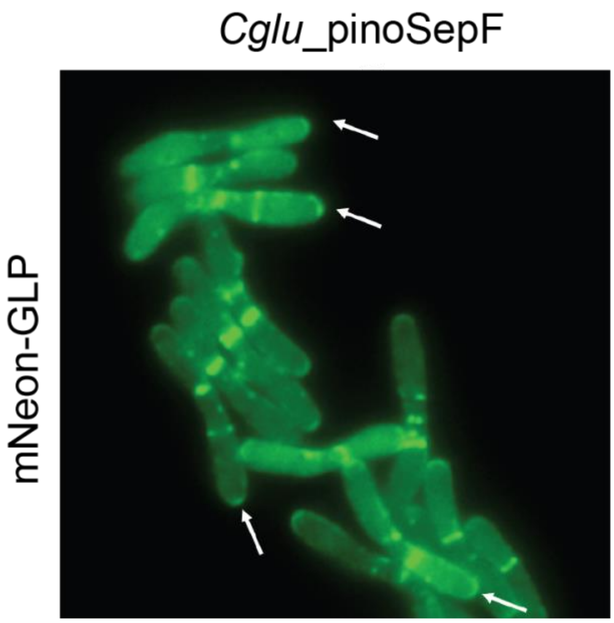

**Figure S8**. Localization of mNeon-GLP in a *Cglu_pinoSepF* background. In the absence of the divisome, GLP can be found in the poles (arrows). SepF was depleted by adding myo-inositol to the culture medium (Sogues et al, 2020).

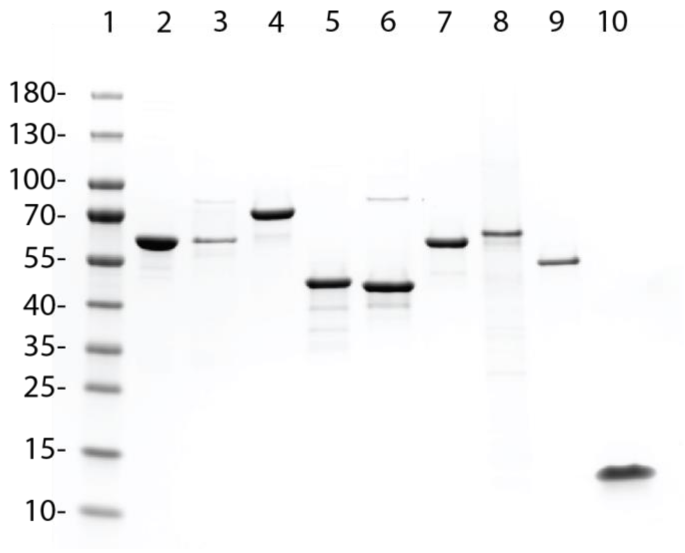

**Figure S9.** SDS-PAGE on a 4-20% polyacrylamide gel of all purified recombinant proteins used in this work. The molecular weight markers (kDa) are indicated on the gel. Lane 1: Molecular weight ladder; lane 2: His-Sumo-GLP (56.3 kDa); lane 3: His-Sumo-GLP$_{\Delta loop}$ (55.1 kDa); lane 4: His-Sumo-FtsZ (59.2 kDa); lane 5: GLP (44.2 kDa); lane 6: GLP$_{\Delta loop}$ (43 kDa); lane 7: FtsZ (47.2 kDa); lane 8: His-GLPR (40.7 kDa); lane 9: Wag31 (38.7 kDa); lane 10: Wag31$_{1\text{-}61}$ (7.1 kDa).

Corresponds to Figure 4d

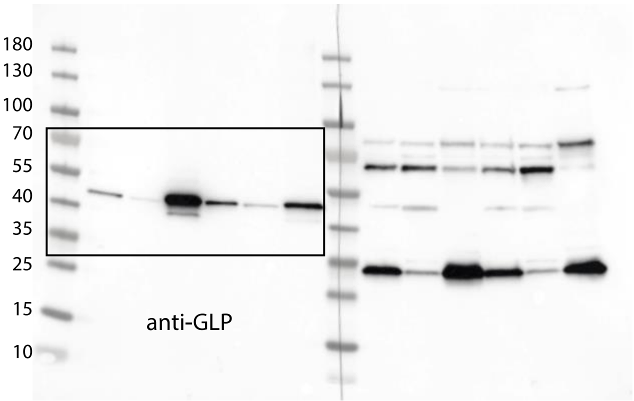

Corresponds to Figure 5h

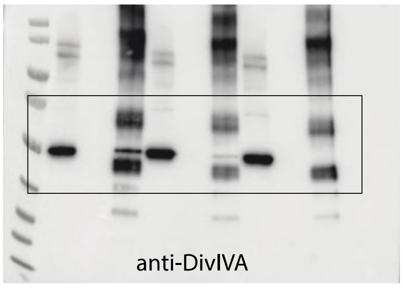

Corresponds to Figure 5h

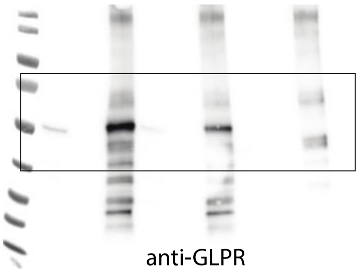

**Figure S10. Uncropped images.** Full uncropped Western Blots of all analyses shown in this work. The boxes correspond to the crops used in the named figures. In Figure 5h, background signals in elution fractions corresponds to the anti-GLPR rabbit antibodies used for the co-IP, which are recognized by the secondary anti-rabbit antibodies used to reveal the Western Blot.

**Table S1. Interactomic data.** (a) SepF interactome, (b) differential interactome $SepF_{K125E/F131A}$ vs SepF (or derivates), (c) GLP interactome in wild-type *Cglu* background, (d) GLP interactome in *Cglu_Δglpr* background.

See Excel Table submitted

**Table S2**. Crystallographic data collection and refinement statistics.

| **Data collection** | **GLP $Cl_4K_2Pt$ derivative** | **GLP** | **GLP FtsZ-CTD** |
|---|---|---|---|
| Synchrotron Beamline | SOLEIL Proxima 1 | SOLEIL Proxima 2A | SOLEIL Proxima 2A |
| Wavelength (Å) | 1.0720 | 0.9801 | 0.9801 |
| Space group | $P4_32_12$ | $P4_32_12$ | $P4_32_12$ |
| Cell dimensions *a*, *b*, *c* (Å) | 94.46, 94.46, 230.61 | 94.44, 94.44, 230.56 | 95.58, 95.58, 229.52 |
| Resolution (Å) | 49.2 – 2.35 (2.41 – 2.35) * | 49.2 – 2.14 (2.19 – 2.14) | 49.2 – 2.68 (2.81 – 2.68) |
| *R*pim | 0.018 (0.186) | 0.023 (0.352) | 0.050 (0.296) |
| *I* / *s*(*I*) | 26.8 (3.9) | 21.1 (2.2) | 14.6 (2.7) |
| Completeness (%) | 99.7 (96.8) | 99.7 (96.2) | 99.6 (97.1) |
| CC(1/2) | 1.0 (0.92) | 1.0 (0.777) | 0.997 (0.800) |
| Multiplicity | 26.8 (26.6) | 26.4 (25.0) | 9.6 (9.7) |
| Total observations | 1197065 | 1551838 | 251139 |
| Unique observations | 44612 (3156) | 58764 (4332) | 30746 (3905) |
| **Refinement** | | | |
| Resolution (Å) | | 2.14 | 2.68 |
| No. reflections | | | |
| *R*work/ *R*free (%) | | 0.191 / 0.223 | 0.190 / 0.239 |
| No. atoms | | | |
| Protein | | 5823 | 5959 |
| Ligands/ions | | - | - |
| Solvent | | 457 | 234 |
| Average B-factors (Å$^2$) | | | |
| Protein | | 49 | 42 |
| Ligand/ions | | - | - |
| Solvent | | 57 | 50 |
| R.m.s deviations | | | |
| Bond lengths (Å) | | 0.007 | 0.004 |
| Bond angles (º) | | 0.876 | 0.663 |
| Ramachandran favored (%) | | 98.0 | 98.6 |
| Ramachandran outliers (%) | | 0 | 0.25 |
| **PDB code** | | 8BVE | 8BVF |

*Values in parenthesis correspond to the highest resolution shell.

**Table S3. Taxonomic sampling of Actinobacteria and protein identifiers of GLP, GLPR and MoeA paralogs**. Protein identifiers correspond to the NCBI GenBank database. “NA” indicates that the protein was not identified in the corresponding genome.

See Excel Table submitted

**Table S4. Taxonomic sampling of Bacteria and protein identifiers of MoeA paralogs**. Protein identifiers correspond to the NCBI GenBank database. “NA” indicates that the protein was not identified in the corresponding genome.

See Excel Table submitted

### Table S5. Plasmids and strains used in this study.

| Strains | Characteristics | Reference |
|---|---|---|
| ***E. coli*** | | |
| DH5*α* | F- endA1 Φ80dlacZΔM15 Δ(lacZYA-argF)U169 recA1 relA1 hsdR17(rK–mK+) deoR supE44 thi-1 gyrA96 phoA λ–; strain used for general cloning procedures | [1] |
| CopyCutter EPI400 | F- mcrA Δ(mrr-hsdRMS-mcrBC) Φ80dlacZΔM15 ΔlacX74 recA1 endA1 araD139 Δ(ara, leu)7697 galU galK λ- rpsL (StrR) nupG trfA tonA pcnB4 dhfr; strain used for general cloning procedures | [2] |
| BL21(DE) | F- ompT hsdSB(rB–mB–) gal dcm (DE3); host for protein production | [3] |
| ***C. glutamicum*** | | |
| ATCC 13032 | Biotin-auxotrophic wild type | [4] |
| ΔGLP | *C. glutamicum* ATCC13032 derivative with chromosomic deletion of GLP | This work |
| ΔGLPR | *C. glutamicum* ATCC13032 derivative with chromosomic deletion of GLPR | This work |
| $P_{ino}$-sepF | *myo*-inositol dependent *sepF* silencing strain. ATCC 13032 with insertion of a terminator and *Pino* promoter to silence *sepF* *(cg2363)* expression. Repressible in the presence of *myo*-inositol | [5] |

| Plasmids for *C. glutamicum* knock out generation | | Reference |
|---|---|---|
| *pK19mobsacB* | KanaR; plasmid for allelic exchange in *C. glutamicum*; (pK18 oriVEc, sacB, lacZα) | [6] |
| pk19-ΔGLP | KanaR; pK19mobsacB derivative for GLP chromosomic deletion | This work |
| pk19-ΔGLPR | KanaR; pK19mobsacB derivative for GLPR chromosomic deletion | This work |

| Plasmids for recombinant protein expression in *E. coli* | | Reference |
|---|---|---|
| pET-SUMO-FtsZ | KanaR; pET derivate for *C. glutamicum* FtsZ recombinant expression containing a N-terminal His-tag followed by a SUMO protease cleavage site | [5] |
| pET-SUMO-GLP | KanaR; pET derivate for *C. glutamicum* GLP recombinant expression containing a N-terminal His-tag followed by a SUMO protease cleavage site | This work |
| pET-SUMO-$GLP_{\Delta Loop}$ | KanaR; pET derivate for *C. glutamicum* $GLP_{\Delta Loop}$ mutant recombinant expression containing a N-terminal His-tag followed by a SUMO protease cleavage site | This work |
| pET-SUMO-Wag31 | KanaR; pET derivate for *C. glutamicum* Wag31 recombinant expression containing a N-terminal His-tag followed by a SUMO protease cleavage site | This work |
| pET-His-TEV-Wag31 | KanaR; pET derivate for *C. glutamicum* Wag31 recombinant expression containing a N-terminal His-tag followed by a TEV protease cleavage site | This work |
| pET-His-TEV-$Wag31_{1\text{-}61}$ | KanaR; pET derivate for *C. glutamicum* N-teminal DivIVA domain of Wag31 (1-61) recombinant expression containing a N-terminal His-tag followed by a TEV protease cleavage site | This work |
| pET-SUMO-$GLPR_{IDR1}$ | KanaR; pET derivate for the recombinant expression of GLPR IDR1 domain (24-214) containing a N-terminal His-tag followed by a SUMO protease cleavage site. | This work |

| | | |
|---|---|---|
| pET-His-GLPR | AmpR; pET derivate for the recombinant expression of GLPR containing a N-terminal His-tag followed by a TEV protease cleavage site | This work |
| pET-GLPR-His | KanaR; pET derivate for the recombinant expression of GLPR containing a C-terminal His-tag. | This work |

| ***Plasmids for recombinant protein expression in C. glutamicum.*** | | **Reference** |
|---|---|---|
| *pTGR5* | KanaR; *E. coli/C. glutamicum* shuttle vector for regulated gene expression of EGFP under control of tac promoter (Ptac lacI ColE1 oriVEc pGA1 oriVCg) | [7] |
| *pUMS3* | KanaR; pTGR5 derivative in which *Ptac* was exchanged by *PgntK* promoter to control the expression of the EGFP protein | [5] |
| pUMS3-PgntK | KanaR; pUMS3 derivative containing *PgntK* promoter (empty plasmid) | This work |
| *pUMS3-sepF-scarlet* | KanaR; pUMS3 derivative for expression of SepF-Scarlet under control of *PgntK* promoter | [5] |
| *pUMS3-sepF*$_{K125/F131A}$ *-scarlet* | KanaR; pUMS3 derivative for expression of SepF$_{K125E-F131A}$-Scarlet mutant under control of *PgntK* promoter | [5] |
| *pUMS3-Scarlet-I* | KanaR; pUMS3 derivative for expression of Scarlet-I fluorescent protein under control of *PgntK* promoter | [5] |
| pUMS3-GLP | KanaR; pUMS3 derivative for expression of GLP under control of PgntK promoter | This work |
| pUMS3-mNeon-GLP | KanaR; pUMS3 derivative for expression of mNeonGreen-GLP under control of PgntK promoter | This work |
| pUMS3-mNeon-GLP$_{\Delta Loop}$ | KanaR; pUMS3 derivative for expression of mNeonGreen-GLP$_{\Delta Loop}$ under control of PgntK promoter | This work |
| pUMS3-GLPR-mNeon | KanaR; pUMS3 derivative for expression of GLPR-mNeonGreen under control of PgntK promoter | This work |
| pUMS3-GLPR | KanaR; pUMS3 derivative for expression of GLPR under control of PgntK promoter | This work |
| pUMS3-GLPR$_{\Delta IDR2}$ (1-266) | KanaR; pUMS3 derivative for expression of GLPR with a deletion of the C-terminal IDR2 region under control of PgntK promoter | This work |
| pUMS3-mNeon-MoeA1 | KanaR; pUMS3 derivative for expression of mNeonGreen-MoeA1 under control of *PgntK* promoter | This work |
| pUMS3-mNeon-MoeA3 | KanaR; pUMS3 derivative for expression of mNeonGreen-MoeA3 under control of *PgntK* promoter | This work |
| pUMS3-Wag31-mNeon | KanaR; pUMS3 derivative for expression of Wag31-mNeonGreen under control of *PgntK* promoter | This work |

**Table S6. Oligonucleotide primers used in this study.**

| Oligonucleotide | Sequence 5' -->3' and properties[a] |
|---|---|
| **Plasmids for *C. glutamicum* knock out generation** | |
| **pk19-ΔGLP** | |
| OligoAS_p68 | TGAGCGGATAACAATTTCAC |
| OligoAS_p69 | CAATTCCACACAACATACG |
| OligoAS_p207 | **TGTTGTGTGGAATTG**CTTGCACACTTTGAGCGTTCTTC |
| OligoAS_p208 | **TTACGCATCGAATA**CGGACCTCCTAATCGGAAC |
| OligoAS_p209 | **CCGTA**TTCGATGCGTAATGCACCGTC |
| OligoAS_p210 | **AATTGTTATCCGCTCA**GCAGATTTCTAATGAGATAGCCTTCTGG |
| **pk19-ΔGLPR** | |
| OligoAS_p68 | TGAGCGGATAACAATTTCAC |
| OligoAS_p69 | CAATTCCACACAACATACG |
| OligoAS_p216 | **TGTTGTGTGGAATTG**ATAAGGAATTCCTCAAGCCCGT |
| OligoAS_p217 | **CTTACCCTGTGG**CTAACCTTCCCGTACGGGTGC |
| OligoAS_p218 | **GTTAG**CCACAGGGTAAGGTTTCGGACTA |
| OligoAS_p219 | **AATTGTTATCCGCTCA**CTGGGAAGTCATACTTCTTGTCCCAC |
| **Oligonucleotides for KO screening** | |
| OligoAS_p211 | CCTATCGATGAGCACGTGAA |
| OligoAS_p212 | ATGCTTCTCCAGCCTTAGCA |
| OligoAS_p220 | GCACCTATTGGCAGGATTGT |
| OligoAS_p221 | TTCCCACATAACCGAGGAAC |

| **Plasmids for recombinant protein expression in *E. coli*** | |
|---|---|
| **pET-SUMO-GLP** | |
| OligoAS_P1 | **AGATCCGGCTGCTAAC**AAAGCCCGAAAG |
| OligoAS_P2 | **GAGGCTCACCGCGAA**CAGATTGGTGGC |
| OligoAS_p169 | **CGAACAGATTGGTGGC**GTGCGATCAGTCGAGCAAC |
| OligoAS_p170 | **GTTAGCAGCCGGATCT**CTATCGACCTTGGGCAAGGAA |
| **pET-SUMO-GLP$_{\Delta Loop}$** | |
| OligoMM_372 | **CATCAGGTCCAGGCTCGGTGTGGAGGCTTTG**GGTGGTGCAACGGGCGCACCATCGCACCT ATTGGCA |
| OligoMM_373 | **CAAAGCCTCCACACCGAGCCTGGACCTGATG**AAACCTTTTCGACCCGCCACAGACACAAC |
